# Human giant GTPase GVIN1 forms an antimicrobial coatomer around the intracellular bacterial pathogen *Burkholderia thailandensis*

**DOI:** 10.1101/2025.03.24.645074

**Authors:** Weilun Guo, Shruti S Apte, Mary S Dickinson, So Young Kim, Miriam Kutsch, Jörn Coers

## Abstract

Several human pathogens exploit the kinetic forces generated by polymerizing actin to power their intracellular motility. Human cell-autonomous immune responses activated by the cytokine interferon-gamma (IFNγ) interfere with such microbial actin-based motility, yet the underlying molecular mechanisms are poorly defined. Here, we identify the IFNγ-inducible human giant GTPases GVIN1 as a novel host defense protein that blocks the bacterial pathogen *Burkholderia thailandensis* from high-jacking the host’s actin polymerization machinery. We found that GVIN1 proteins form a coatomer around cytosolic bacteria and prevent *Burkholderia* from establishing force-generating actin comet tails. Coatomers formed by a second IFNγ-inducible GTPase, human guanylate binding protein 1 (GBP1), constitute a GVIN1-independent but mechanistically related anti-motility pathway. We show that coating with either GVIN1 or GBP1 displaces the *Burkholderia* outer membrane protein BimA, an actin nucleator that is essential for actin tail formation. Both GVIN1 and GBP1 coatomers require additional IFNγ-inducible co-factors to disrupt the membrane localization of BimA, demonstrating the existence of two parallel-acting IFNγ-inducible defense modules that evolved to target a virulence trait critical for the pathogenesis of numerous bacterial infectious agents.

## INTRODUCTION

Actin-based bacterial motility is a key mechanism by which host cytosol-invading bacterial pathogens like *Listeria, Shigella,* and *Burkholderia* move within and between infected cells. By recruiting and polymerizing host actin filaments at the surface of one bacterial pole, these bacteria form actin “comet tails” that propel the bacteria forward, enabling them to move rapidly through the infected cell and directly into neighboring cells (*1, 2*). This dissemination mechanism enables pathogens to spread within cell layers while avoiding extracellular immunity. Intracellular cell-autonomous host defenses that target actin motility are thus critical for immune protection, yet our understanding of these defenses is limited.

The general mechanisms by which intra-cytosolic bacterial pathogens hijack the host cell’s actin polymerization machinery are well characterized. For example, in *Shigella* the outer membrane autotransporter IcsA is localized to one bacterial pole and recruits the host protein N-WASP, an actin nucleation promotion factor. N-WASP in turn recruits the actin polymerization initiation Arp2/3 complex to launch actin filament formation (*3–7*). Like IcsA in *Shigella,* the *Burkholderia* outer membrane protein BimA is localized to one bacterial pole and is essential for actin-based motility (*8, 9*). BimA of *Burkholderia thailandensis* is an N-WASP mimic that directly recruits Arp2/3 to the bacterial pole to initiate actin polymerization (*10*).

The polar localization of bacterial proteins like IcsA and BimA is essential for their function and tightly regulated through multiple mechanisms. Specifically, the function of IcsA is dependent on lipopolysaccharide (LPS), the main building block of the outer leaflet of Gram-negative bacterial outer membranes. LPS consists of its membrane anchor lipid A, an inner core oligosaccharide, and the outer core polysaccharide O antigen. The outwards-facing thick O antigen layer provides a chemo-physical barrier that protects bacteria against antimicrobial molecules (*11*). Additionally, O antigen enhances bacterial outer membrane stiffness (*12*). Accordingly, loss of O antigen in ‘rough mutant’ bacteria results in increased outer membrane fluidity and enhanced diffusion of outer membrane proteins (*12*). The functional importance of this O antigen-mediated membrane stiffness is illustrated by the enhanced diffusion of IcsA proteins away from the bacterial pole and the consequential loss of actin-based motility in *Shigella flexneri* rough mutants (*13–15*). Whether O antigen is also required for the function of the actin nucleator BimA of *Burkholderia* has not been previously reported.

Actin-based cell-to-cell movement not only facilitates bacterial spread within tissues, but also helps pathogens avoid extracellular immunity. To control infections with pathogens that largely remain hidden intracellularly, host cells are equipped with cell-autonomous immune responses. These cell-autonomous defense programs are executed by a range of immune effector proteins, many of which are expressed in response to proinflammatory cytokine signaling. Arguably, the most potent inducer of antibacterial cell-autonomous immunity is the lymphocyte-derived cytokine IFNγ (*16*). Among the hundreds of IFNγ-inducible genes are those encoding members of GTPase families that include ‘Immunity-Related GTPases’ (IRGs), giant GTPases annotated as ‘GTPases, Very large Interferon-inducible’ (GVINs), and ‘Guanylate Binding Proteins’ (GBPs). Whereas GBPs and other interferon-inducible GTPases families have proven antimicrobial activities, no biological function has previously been reported for GVIN proteins (*17*).

The most highly expressed human GBP family member is GBP1, an LPS-binding protein that can form coatomers on the surface of Gram-negative bacteria including *Shigella* and *Burkholderia* (*18–20*). Once embedded onto the bacterial outer membrane, the GBP1 coatomer operates akin to a surfactant that disrupts the function of O antigen (*13*). As a result, IcsA is displaced from the bacterial pole of GBP1-coated bacteria (*13*), which consequently lose their ability to form actin tails and to disseminate within epithelial monolayers (*13, 21, 22*). Human GBP1 coatomers also assemble on the surface of *B. thailandensis* (*21*). However, studies conducted in human HeLa epithelial cells indicated that these GBP1 coatomers formed on the surface of *B. thailandensis* fail to disrupt the formation of actin tails. Instead, it was suggested that GBP1-mediated host cell death may inhibit the intercellular spread of *B. thailandensis* (*23*). In contrast to these findings in human HeLa cells, a separate study demonstrated that the mouse ortholog of human GBP1, mouse Gbp2, was capable of directly interfering with *B. thailandensis* actin tail formation in murine primary macrophages (*24*). However, the underlying mechanism by which mouse Gbp2 modulates *Burkholderia*’s actin-based motility was not further explored.

In the current work we demonstrate that some human cell lines block *B. thailandensis* actin tail formation when primed with IFNγ, whereas other human cell lines including commonly used HeLa cervical epithelial cells lack the ability to interfere with *B. thailandensis* actin tail formation. We find that IFNγ priming can activate two independent but functionally related anti-motility pathways that are dysfunctional in HeLa cells but functional in T24 bladder epithelial cells. The first pathway requires the formation of GBP1 coatomers and the expression of additional IFNγ-inducible factors that are ostensibly non-functional or not expressed in HeLa cells. A second anti-motility pathway is dependent on the expression of GVIN1 (GTPase, Very large IFN-Inducible-1), the single human member of the GVIN family of IFN-inducible giant GTPases (*17*). Because automated annotation pipelines flag predicted in-frame stop codons, GVIN1 has long been catalogued as the pseudogene GVINP1 (GTPase, Very large IFN-Inducible Pseudogene-1) by Ensembl, GENCODE, and RefSeq (HGNC 25813; NCBI Gene 387751; OMIM 616121). In contrast to its designation as a pseudogene, we show here that human GVIN1is a functional protein that forms a coatomer surrounding cytosolic *B. thailandensis*. Like GBP1 coatomers, GVIN1 coatomers require additional IFNγ-inducible factors to disrupt the localization of BimA and to interfere with actin tail formation. Our study thus identifies GVIN proteins as a novel class of antimicrobial coatomer-forming proteins.

## RESULTS

### IFNγ priming restricts *B. thailandensis* actin tail formation in a subset of human cell lines

Confirming previous observations (*13, 21, 22*), we found that GBP1 coatomers form on the surface of *S. flexneri* in IFNγ-primed human HeLa and T24 epithelial cell lines, prevent recruitment of Arp2/3, and block bacterial actin-tail formation (Fig. S1A-G), as visualized by phalloidin staining. Although GBP1 coatomers also form on the surface of *B. thailandensis* in IFNγ-primed HeLa cells (Fig. 1A-B), such GBP1 coatomers failed to diminish Arp2/3 recruitment to the bacterial pole (Fig. S1H-I) and had no discernable impact on the percentage of bacteria that formed actin tails (Fig. 1C). These data confirmed that IFNγ-primed HeLa cells block actin tail formation by *S. flexneri* but not by *B. thailandensis*.

**Fig. 1.**
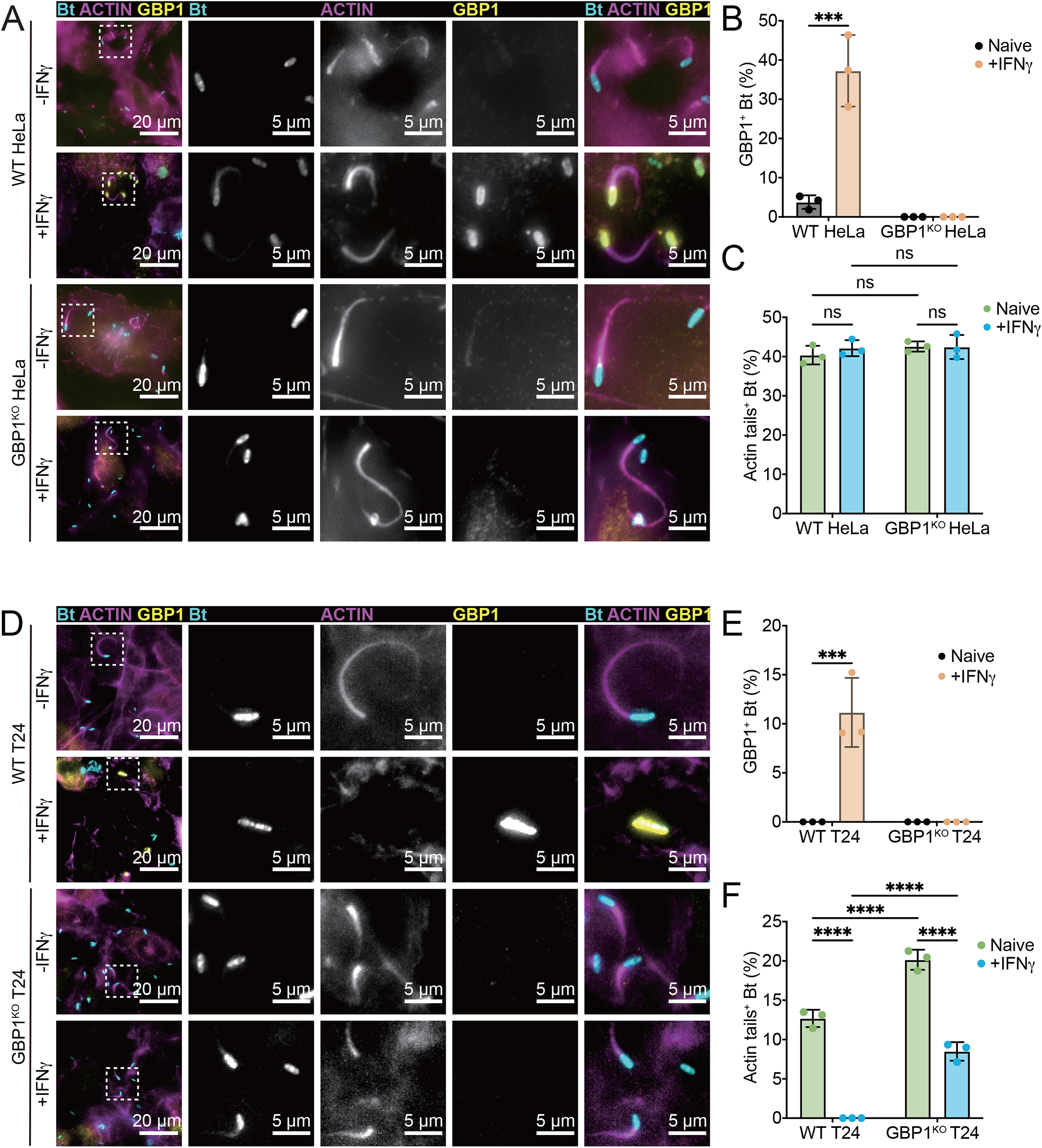
IFNγ-primed T24 cells restrict actin tails formation by *B. thailandensis.* (A-C) Naïve or 200 U/mL IFNγ primed WT or GBP1^KO^ HeLa cells were infected with WT *B. thailandensis* at an MOI of 100. Cells were fixed at 8 hours post infection and stained for GBP1 and actin. Percentages of GBP1-positive *B. thailandensis* (B) and bacteria with actin tails (C) were quantified. (D-F) Naïve or 200 U/mL IFNγ primed WT or GBP1^KO^ T24 cells were infected with WT *B. thailandensis* at an MOI of 100. Cells were fixed at 8 hours post infection and stained for GBP1 and Actin. Percentages of GBP1-positive *B. thailandensis* (E) and bacteria with actin tails (F) were quantified. All bar graphs show 3 independent biological replicates, presented as mean ± standard deviation (SD). Two-way ANOVA with Tukey’s multiple comparison tests were performed, with specific p-values indicated as follows: ***, p < 0.001; ****, p < 0.0001; ns, not significant.

To more broadly test whether any human cell lines exist that could block actin tail formation by *B. thailandensis*, we monitored the effect of IFNγ priming on actin tail formation in a small collection of commonly used human cell lines that included A549 lung epithelial cells, T24 bladder epithelial cells, and differentiated monocytic U937 and THP1 cells. In the case of THP1 monocytes we used CASP4-deficient (CASP4^KO^) cells to avoid the confounding effects of *B. thailandensis*-triggered CASP4-driven pyroptotic cell death. We observed that IFNγ priming significantly diminished actin tail formation by *B. thailandensis* in T24, U937, and THP1-CASP4^KO^ but not in A549 cells (Fig. 1D-F and S2A-C). Deletion of GBP1 in IFNγ-primed epithelial T24 cells partially restored actin tail formation by *B. thailandensis* (Fig. 1D and 1F), indicating that both GBP1-dependent and GBP1-independent pathways impede *B. thailandensis* actin tail formation in human T24 cells. The GBP1-independent pathway is ostensibly inactive against *S. flexneri* based on the observation that the reduction in actin tail formation by *S. flexneri* in IFNγ-primed T24 cells is completely reversed by the deletion of GBP1 (Fig. S1F-G).

Previous work suggested that GBP1 may limit cell-to-cell spread by *B. thailandensis* through the induction of pyroptotic cell death in HeLa cells (*23*). However, such a mechanism is unlikely to be active in T24 cells, because we found that naïve and IFNγ-primed WT and GBP1^KO^ T24 cells displayed comparable levels of cell death as assessed by LDH release assays (Fig. S2D).

### GBP1-mediated inhibition of *B. thailandensis* actin tail formation requires a co-factor that is missing from HeLa cells

Hela cells lack not only the GBP1-independent but also the GBP1-dependent pathway to restrict *B. thailandensis* actin tail formation, despite having GBP1 protein expression levels that are comparable to T24 cells (Fig. 2A). We therefore considered that the loss of the GBP1-dependent pathway in HeLa cells was either due to i) differences in the GBP1 protein sequence between the two cell lines or ii) the absence of an essential co-factor from HeLa cells. To test both hypotheses in one experiment, we engineered GBP1^KO^ T24 and GBP1^KO^ HeLa cells that inducibly expressed the same N-terminally mCherry-tagged GBP1 (mCh-GBP1) fusion protein upon treatment of cells with anhydrotetracycline (aTc) (Fig. 2B). Targeting of mCh-GBP1 to *B. thailandensis* occurred at comparable frequency in GBP1^KO^ T24 and GBP1^KO^ HeLa cells (Fig. S2E-F). We found that mCh-GBP1 overexpression was sufficient to reduce the percentage of actin tail-positive *B. thailandensis* by 2-fold in in unprimed GBP1^KO^ T24 but not in unprimed GBP1^KO^ HeLa cells (Fig. 2C-D). These data suggest that T24 but not HeLa cells express an essential GBP1 co-factor, even in the absence of IFNγ priming. Combining ectopic mCh-GBP1 expression with IFNγ priming led to a near complete elimination of actin tail formation in T24 GBP1^KO^ but had no discernable effect on bacterial actin tail formation in HeLa GBP1^KO^ cells (Fig. 2C-D). IFNγ priming significantly increased the population of mCh-GBP1 coated bacteria that are actin negative (Fig. S2G), further highlighting the synergistic effects between GBP1 and additional IFNγ-inducible host factors. The cooperation between ectopic GBP1 expression and IFNγ priming in T24 GBP1^KO^ cells could be largely explained by the induction of a GBP1-independent host defense pathway in IFNγ-primed T24 cells. However, IFNγ stimulation may additionally enhance GBP1-dependent host defense against *B. thailandensis*, if the putative GBP1 co-factor is itself an IFNγ-inducible gene (ISG).

**Fig. 2.**
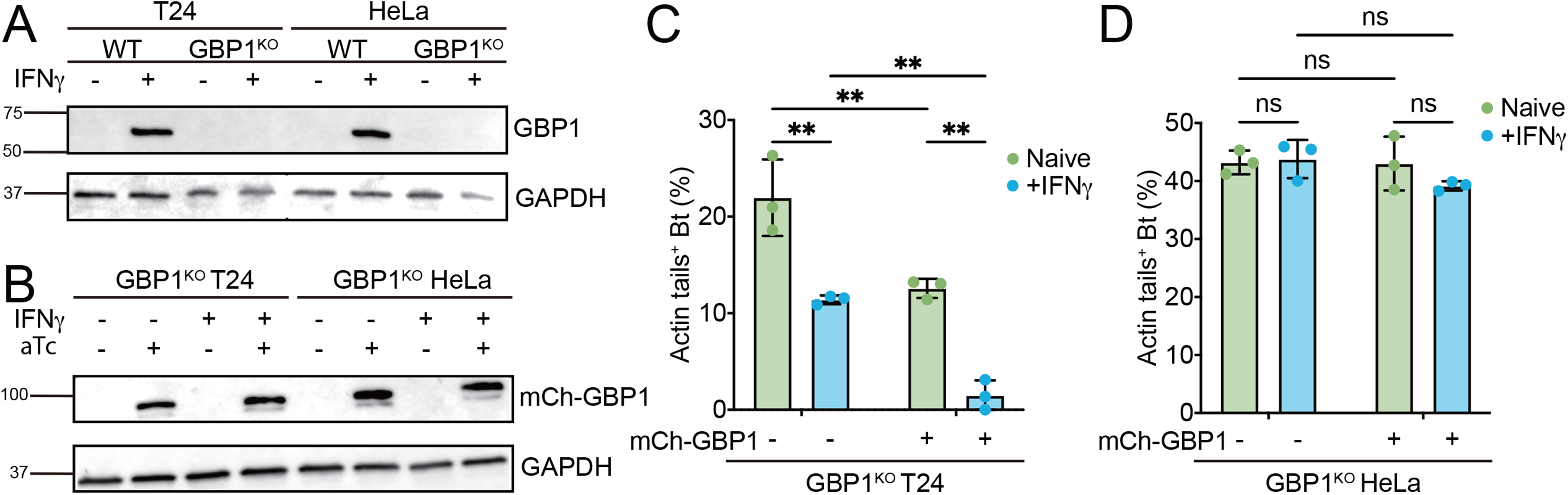
GBP1 requires a co-factor to restrict *B. thailandensis* actin tail formation. (A) Immunoblotting for GBP1 protein expression in WT HeLa, GBP1^KO^ HeLa, WT T24, or GBP1^KO^ T24 cells treated with or without 200 U/ml IFNγ. (B-D) Parental GBP1^KO^ HeLa or GBP1^KO^ T24 cells were transduced with an anhydrotetracycline (aTc)-inducible mCherry-GBP1 lentiviral construct (mCh-GBP1). (B) Immunoblotting for GBP1 in naïve or 200 U/mL IFNγ primed GBP1^KO^ HeLa + mCh-GBP1 or GBP1^KO^ T24 + mCh-GBP1cells treated with or without 1 μg/mL aTc for overnight. (C-D) Naïve or 200 U/mL IFNγ primed GBP1^KO^ T24 + mCh-GBP1 (C) or GBP1^KO^ HeLa + mCh-GBP1 (D) cells were treated with or without 1 μg/mL aTc for overnight, and then infected with WT *B. thailandensis* at an MOI of 100. Cells were fixed at 8 hours post infection and stained for actin. Percentages of *B. thailandensis* with actin tails were quantified. All bar graphs show 3 independent biological replicates, presented as mean ± SD. Two-way ANOVA with Tukey’s multiple comparison tests were performed, with specific p-values indicated as follows: **, p < 0.01; ns, not significant.

### Loss of O antigen renders *B. thailandensis* resistant to GBP1-independent inhibition of actin tail formation

GBP1 coatomer formation is sufficient to block actin formation by *S. flexneri* but not by *B. thailandensis* (Figs. 1, S1, and 2 and (*13, 21, 22*)). The mechanism by which GBP1 coatomers block actin tail formation was previously elucidated for *S. flexneri*: GBP1 coatomers on the surface of *S. flexneri* disrupt the O antigen layer and thereby turn bacteria into quasi-O antigen deficient strains. Because O antigen is essential for the polar localization and function of IcsA (*13–15*), formation of GBP1 coatomers on *S. flexneri* is sufficient to block IcsA-dependent actin tail formation (*13*). To explain why GBP1 coatomer formation on the other hand is insufficient to block actin formation by *B. thailandensis*, we considered two hypotheses: i) the O antigen layer of *B. thailandensis* is impervious to disruption by GBP1 coating or ii) O antigen is dispensable for *B. thailandensis* BimA function. To test the latter hypothesis, we generated loss-of-function mutants in the gene BTH_I1483, which encodes the predicted polysaccharide biosynthetic enzyme WbpM (Fig. 3A). Immunofluorescence staining with *B. thailandensis*-specific anti-LPS antibody and ProQ Emerald 300 LPS gel staining characterized the *wbpM* CmR insertion mutant (henceforth simplified as Δ*wbpM*) as O antigen-deficient (Fig. 3B-C). We found that the O antigen-deficient w*bpM* mutant formed actin tails at frequencies comparable to wildtype (WT) bacteria in naïve T24 cells (Fig. 3D-E), thus demonstrating that O antigen is dispensable for *B. thailandensis* actin-based motility. These findings suggest that BimA remains functional even when the membrane stiffening property of O antigen is disrupted by GBP1 coatomers. Unexpectedly, we also observed that the O antigen deficient *B. thailandensis wbpM* mutant became resistant to IFNγ-mediated inhibition of bacterial actin tail formation in T24 GBP1^KO^ cells (Fig. 3E). The latter finding suggested that the GBP1-independet host defense pathway specifically recognizes the O antigen polysaccharide layer of the bacterial outer membrane.

**Fig. 3.**
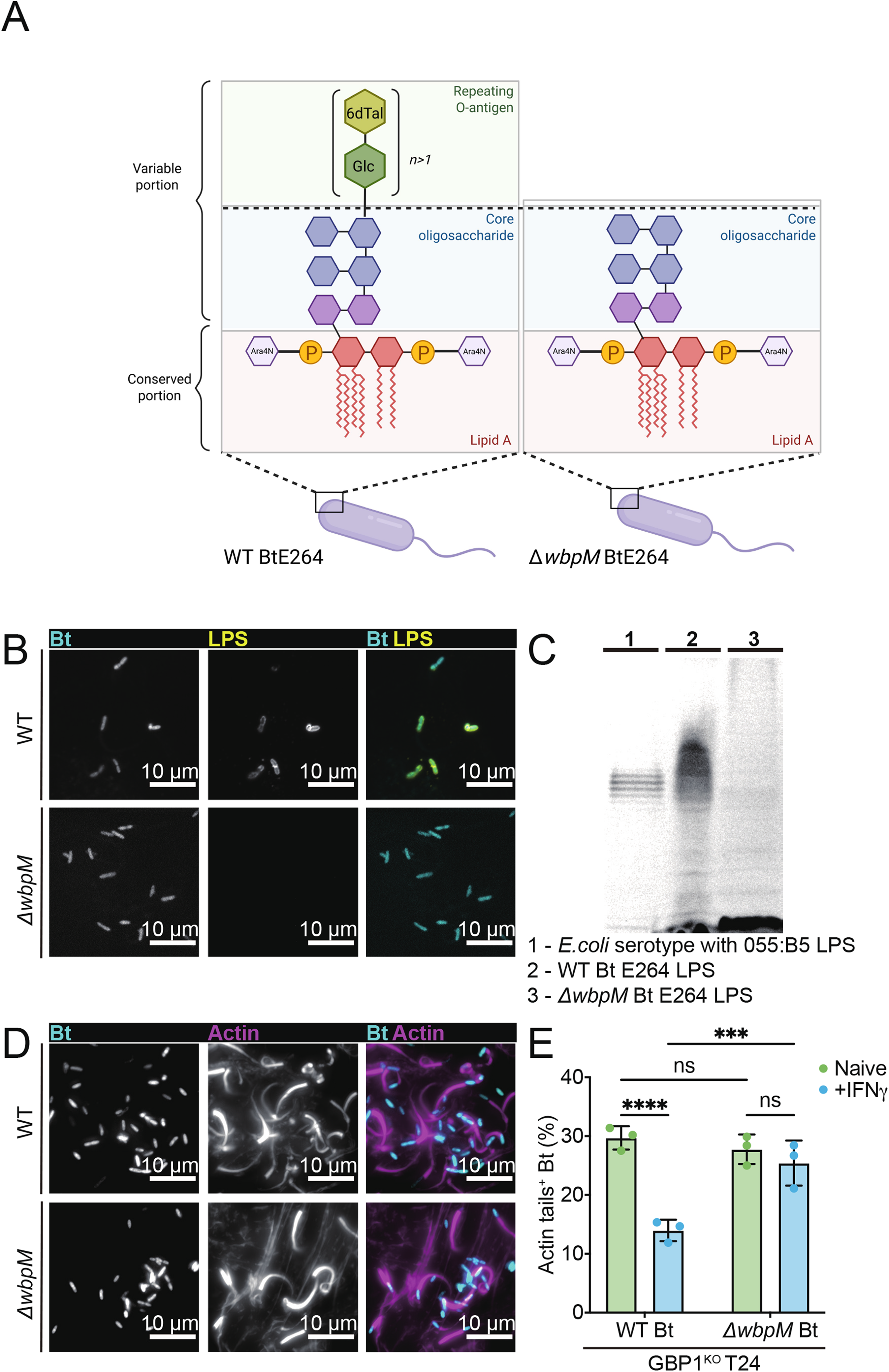
O antigen is dispensable for *B. thailandensis* actin tail formation but promotes ifNγ-inducible host defense. (A) Schematic depiction of WT *B. thailandensis* and *ΔwbpM B. thailandensis* LPS structure. The O antigen is a linear heteropolymer featuring a disaccharide unit in an equimolar ratio of (1→3)-linked 6-deoxy-α-l-talopyranose and β-d-glucopyranose. Adapted from Kenfack et al. 2017 (*60*) (B-E) Naïve GBP1^KO^ T24 cells were infected with WT or *ΔwbpM B. thailandensis* at an MOI of 100. Cells were fixed at 8 hours post infection and stained for *B. thailandensis* LPS (B) and actin (D), and percentages of *B. thailandensis* with actin tails (E) were quantified. (C) SDS-PAGE analysis of LPS from *E.coli* serotype with 055:B5, WT *B. thailandensis* E264, or *ΔwbpM B. thailandensis* E264 with the Pro-Q Emerald 300 stain. All bar graphs show 3 independent biological replicates, presented as mean ± SD. Two-way ANOVA with Tukey’s multiple comparison tests were performed, with specific p-values indicated as follows: ***, p < 0.001; ****, p < 0.0001; ns, not significant.

### A genetic screen identifies GVIN1 as an inhibitor of *B. thailandensis* actin tail formation

We found that IFNγ-inducible pathways interfering with *B. thailandensis* actin tail formation are present in some (T24, U937, THP1) but absent from other (A549 and HeLa) human cell lines (Fig. 1 and S2). We therefore hypothesized that A549 and HeLa cells lack robust expression of specific ISGs that are required for the inhibition of actin tail formation. To identify candidate ISGs, we profiled gene expression in all five cell lines (T24, U937, THP1, A549, and HeLa) under naïve and IFNγ-primed conditions using bulk RNAseq and then used these data to curate a list of candidate genes that we could test for function (Fig. 4A). Based on our RNAseq data we identified 28 protein-coding ISGs that were induced at least 2 log-fold upon IFNγ priming in T24 cells and were expressed at least 2-fold higher in IFNγ-primed T24, U937, and THP1 cells relative to IFNγ-primed A549 or IFNγ-primed HeLa cells (Fig. 4A and S3A-B). To determine whether any of these candidate ISGs were involved in the GBP1-independent pathway, we conducted an arrayed functional genomics ‘mini screen’ in T24 GBP1^KO^ cells. For each candidate gene, individual wells seeded with T24 GBP1^KO^ cells were transfected with four independent siRNA oligonucleotides. We then primed the transfected cells with IFNγ overnight, infected the cells with *B. thailandensis*, and quantified the percentage of bacteria forming actin tails at 8 hours post infection (hpi). The number of cytosolic *B. thailandensis* forming actin tails inside IFNγ-primed T24 GBP1^KO^ cells significantly increased by approximately 2-fold relative to cells treated with siRNA control oligonucleotides for two targeted ISGs, namely *GBP6* (Gene ID: 163351) and *GVINP1* (GTPase, Very large interferon inducible pseudogene 1; Gene ID: 387751) (Fig. 4B and S3C). From hereon we will refer to *GVINP1* as *GVIN1*, as we will provide evidence for it to be a functional gene and not a pseudogene.

**Fig. 4.**
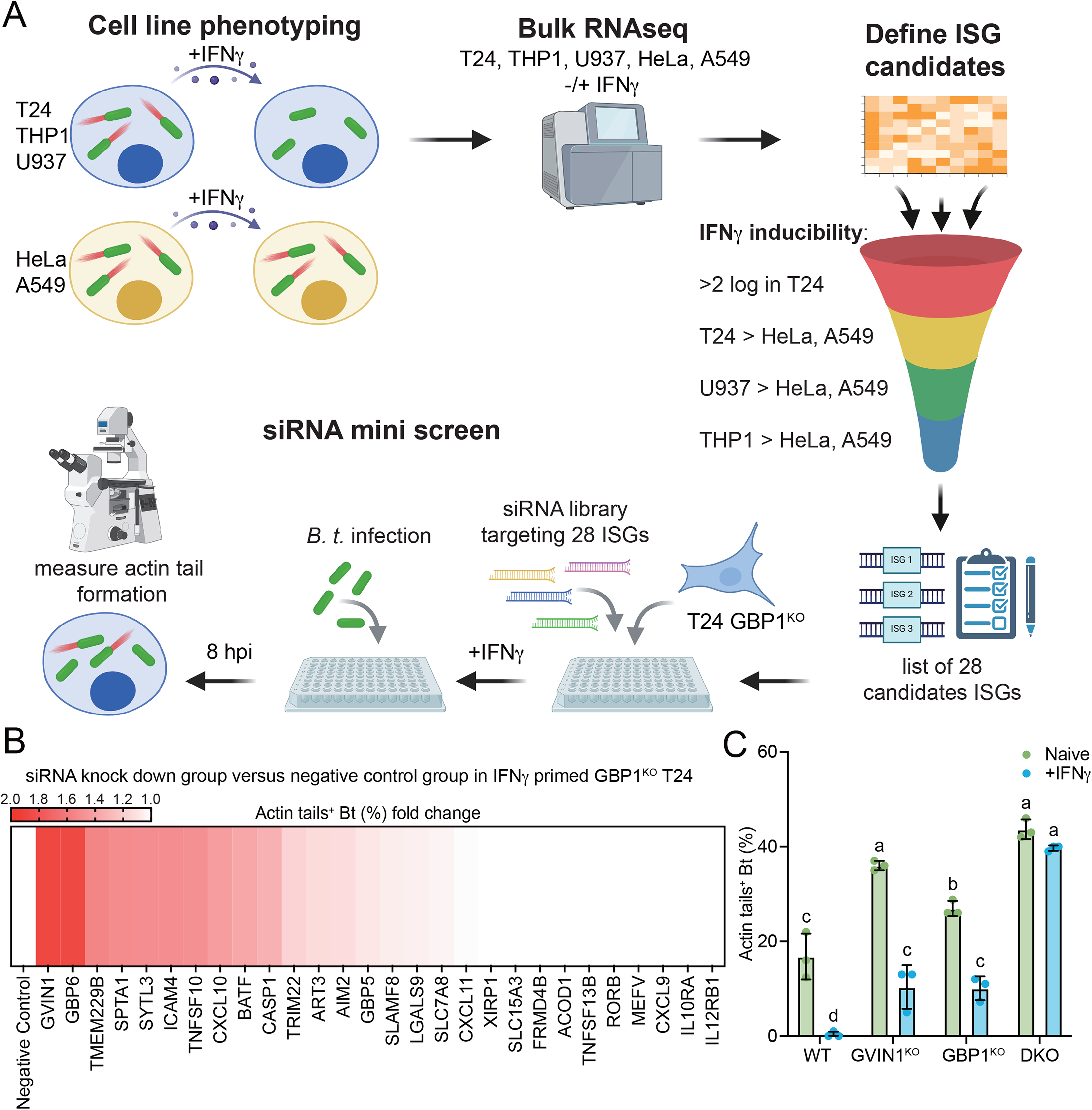
A genetic screen identifies GVIN1 as an inhibitor of *B. thailandensis* actin tail formation. (A) Schematic depiction of the strategy for the siRNA mini screen. (B) 5 pmol of specific gene’s siRNA transfected and 200 U/mL IFNγ primed GBP1^KO^ T24 cells were infected with WT *B. thailandensis* at an MOI of 100. Cells were fixed at 8 hours post infection and stained for actin. Percentages of *B. thailandensis* with actin tails were quantified and showed as fold changes of %Actin-tail^+^ *B. thailandensis* in the specific gene’s siRNA knock down versus in the negative control siRNA transfected GBP1^KO^ T24 were shown in the heat map. (C) Naïve or 200 U/mL IFNγ primed WT, GBP1^KO^, GVIN^KO^, or GBP1^KO^ GVIN^KO^ DKO T24 cells were infected with WT *B. thailandensis* at an MOI of 100. Cells were fixed at 8 hours post infection and stained for actin. Percentages of *B. thailandensis* with actin tails were quantified. All bar graphs show 3 independent biological replicates, presented as mean ± SD. Two-way ANOVA with Tukey’s multiple comparison tests were performed, with specific p-values indicated as follows: Different letters above columns represent there is significant difference, same letter above columns represents not significant.

To independently validate whether GBP6 and GVIN1 play any role in limiting actin tail formation by *B. thailandensis*, we generated *GBP6* and *GVIN1* deletions in T24 cells using CRISPR technology (Fig. S4A-B). We found that IFNγ-primed T24 GBP6^KO^ cells restricted *B. thailandensis* actin tail formations at rates that were comparable to IFNγ-primed wild type T24 cells (Fig. S4C). Furthermore, GBP6 protein expression in naïve or IFNγ-primed T24 cells was below the level of detection by Western blotting (Fig. S4D). Collectively, these data suggested that GBP6 was a false-positive hit from our screen and accordingly was not investigated further. Additionally, the possibility of false-negatives due to variable knockout efficiencies cannot be excluded.

In contrast to GBP6^KO^ cells, GVIN1^KO^ cells allowed for increased rates of actin tail formation by *B. thailandensis* and showed incomplete restriction of actin tail formation upon IFNγ priming, thus mimicking the phenotype of GBP1^KO^ cells (Fig. 4C). Simultaneous deletion of both GVIN1 and GBP1 completely abolished the ability of IFNγ-primed T24 cells to limit actin tail formation by *B. thailandensis* (Fig. 4C). Despite the dramatic reduction in actin tail formation, IFNγ priming of WT T24 cells did not measurably impede cell-to-cell spread by *B. thailandensis,* as assessed by plaque assays (Fig. S5A). A likely explanation for this counterintuitive finding is that *B. thailandensis* can spread from cell to cell independent of actin-drive propulsion using flagellar motility (*25*).

Whereas IFNγ stimulation in T24 eliminated nearly all detectable actin tail formation (Fig. 4C), its effect on *B. thailandensis* viability as measured by colony forming units (CFU) counts was only modest (Fig. S5B). The IFNγ-driven reduction in bacterial burden was dependent on both GBP1 and GVIN1, as bacterial numbers were restored equally in either GBP1^KO^ or GVIN1^KO^ T24 cells (Fig. S5B). Together, these data indicate that GBP1 and GVIN1 are critical players in the execution of two host defense pathways targeting *B. thailandensis.* Our observations indicate that GBP1 and GVIN1 interfere with actin tail formation independently from each other but appear to work cooperatively to reduce bacterial burden.

### GVIN1 coatomers require IFN**γ**-inducible co-factors to block actin tail formation

The *GVIN1* gene is intronless and expresses a transcript that is approximately 8,700 nucleotides in length. The GVIN1 mRNA contains three potential open reading frames (ORFs). The longest uninterrupted ORF is 5,424 nucleotides in length and predicted to encode a protein of 1,808 amino acids with an expected molecular weight of 209 kDa (Fig. 5A), hereinafter simply referred to as GVIN1 protein. To gain insights into the potential function of GVIN1, we employed the protein structure prediction software AlphaFold3 (*26, 27*) and the structural alignment tool Foldseek (*28*). The predicted structure features an N-terminal domain that is connected to a Large GTPase (LG) domain via a flexible, mostly unstructured linker, followed by helical linker domain, a second LG domain, and a C-terminal domain (Fig. 5A-B and S6A-B). Because related GTPases of the GBP family form dimers upon GTP-binding and-hydrolysis (*18*), we also modeled GVIN1 dimerization in AlphaFold3. Our predictions suggest that GVIN1 forms dimers via its LG domains (Fig. S6C-E), which is in line with dimerization of Mx proteins, IRGs, and GBPs (*29*).

**Fig. 5.**
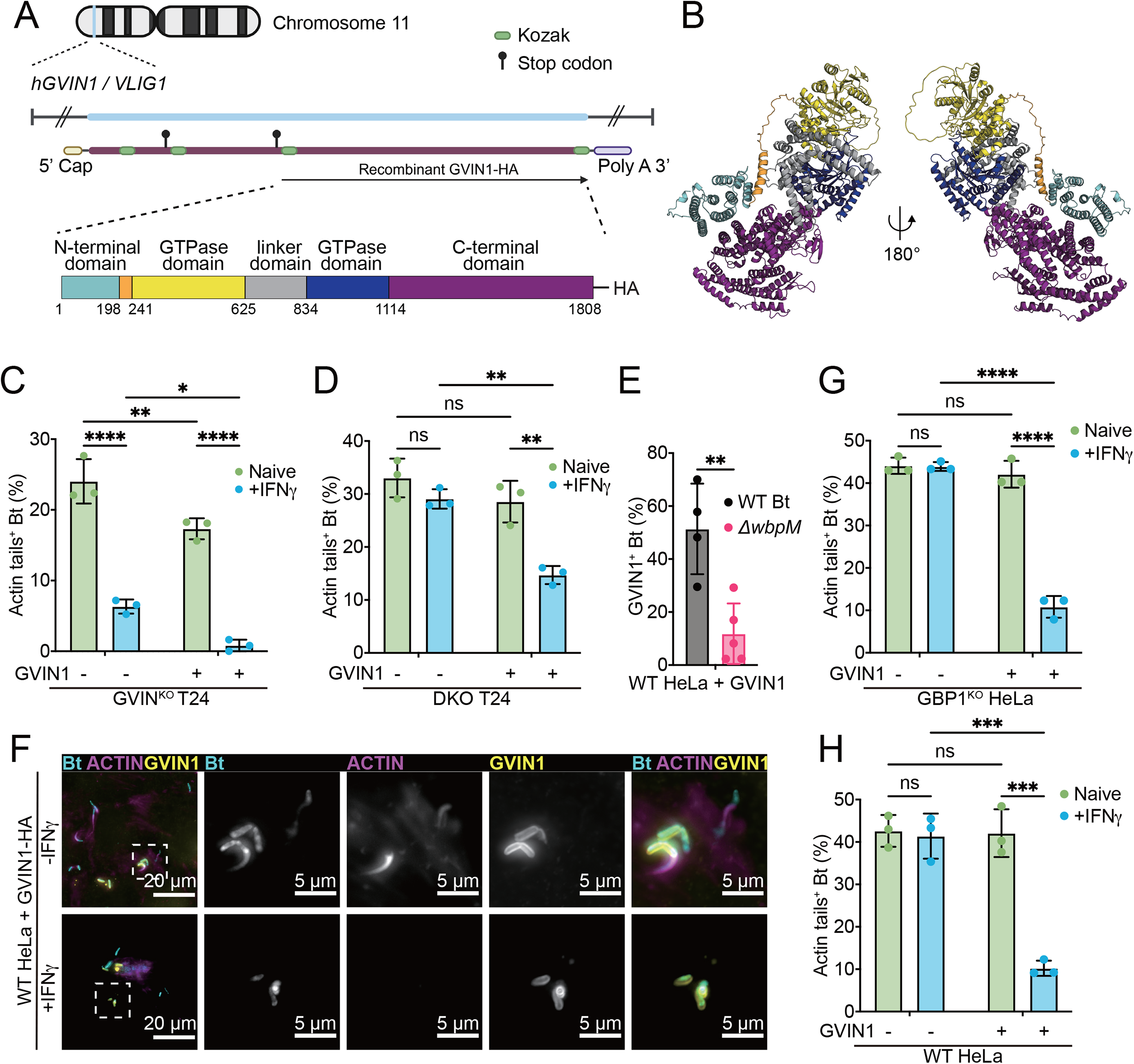
GVIN1 restricts *B. thailandensis* actin tail formation in an ifNγ dependent but GBP1 independent manner. (A) Schematic depiction of the GVIN1 DNA, GVIN1 mRNA and the recombinant GVIN1 protein (C-terminal-HA-tagged protein translated from GVIN1 longest ORF). The domains were assigned by Foldseek of the predicted AlphaFold structure, the numbers reflect the position of the amino acid in the predicted protein sequence, starting with the start amino acid methionine of the longest ORF of GVIN1. (B) 3D structure model of the GVIN1 longest ORF, predicted by AlphaFold3. (C-H) Parental WT HeLa, GBP1^KO^ HeLa, GVIN1^KO^ T24, or GBP1^KO^ GVIN^KO^ DKO T24 cells were transduced with aTc-inducible GVIN1-HA lentiviral construct. IFNγ primed and/or 1 μg/mL aTc overnight-treated WT HeLa + GVIN1-HA (E), GBP1^KO^ HeLa + GVIN1-HA, GVIN1^KO^ T24 + GVIN1-HA, or GBP1^KO^ GVIN^KO^ DKO T24 + GVIN1-HA were infected with WT *B. thailandensis* at an MOI of 100. Cells were fixed at 8 hours post infection and stained for actin. Percentage of actin-tail-positive *B. thailandensis* were quantified (C, D, G, H). (F) Naïve or 200 U/mL IFNγ primed and/or 1 μg/mL aTc overnight-treated WT HeLa + GVIN1-HA cells were infected with WT or *ΔwbpM B. thailandensis* at an MOI of 100. Cells were fixed at 8 hours post infection and stained for GVIN1-HA. Percentages of GVIN1-HA positive *B. thailandensis* were quantified. All bar graphs show 3 independent biological replicates, presented as mean ± SD. Two-way ANOVA with Tukey’s multiple comparison tests were performed for (C), (D), (G), and (H); and unpaired t-test was performed for (F). Specific p-values indicated as follows: *, p < 0.05; **, p < 0.01; ***, p < 0.001; ****, p < 0.0001; ns, not significant.

Dimeric GBP1 proteins insert into Gram-negative bacterial outer membranes to form coatomers and execute host defense (*13, 30–32*). To assess whether GVIN1 could operate in a similar fashion, we expressed a C-terminally HA-tagged GVIN1 protein under control of an aTc-inducible promoter in cell culture. Ectopically expressed GVIN1-HA (henceforth referred to simply as GVIN1) was detectable by Western blotting at approximately its predicted size of 209 kDa (Fig. S7A). Importantly, we found that expression of GVIN1 in T24 GVIN1^KO^ restored IFNγ-dependent restriction of *B. thailandensis* actin tail formation to levels comparable to WT cells (Fig. 5C). Ectopic expression of GVIN1 in T24 GVIN1/GBP1^DKO^ partially restored IFNγ-dependent restriction of actin tail formation comparable to the phenotype of GBP1^KO^ cells (Fig. 5D). These complementation experiments confirmed that loss of functional GVIN1 is responsible for elevated levels of *B. thailandensis* actin tail formation in GVIN1^KO^ cells.

Next, we monitored the subcellular localization of ectopically expressed GVIN1 in *B. thailandensis* infected HeLa cells and observed that at 8 hpi approximately 50% of intracellular bacteria were encased by GVIN1 (Fig. 5E-F). This encapsulation morphologically resembled coatomers formed by GBP1 on *B. thailandensis* (Fig. 1A-B and (*21*)). Actin tail formation in IFNγ-primed T24 cells was largely absent from GVIN1-enwrapped *B. thailandensis* (Fig. S7B), indicating that the GVIN1 coatomer is required for GVIN1-driven inhibition of actin tail formation. Dual immunostaining for GBP1 and GVIN1 revealed that most targeted bacteria were encased either within a GBP1 or a GVIN1 coatomer (Fig. S7C). However, a rare subpopulation of GBP1-/GVIN1-double positive bacteria was detectable in these IFNγ-primed cells (Fig. S7C-D), indicating that the two types of coatomers are generally distinct but not strictly mutually exclusive.

We next investigated possible bacterial factors impacting GVIN1 coatomer formation. Actin tail formation itself was dispensable for GVIN1 recruitment, as evidenced by the comparable rates of GVIN1 coatomer formation between WT and Δ*bimA B. thailandensis* (Fig. S8A). However, GVIN1 coatomers formed only infrequently on the surface of the O antigen deficient Δ*wbpm B. thailandensis* mutant (Fig. 5E, S8B), thus indicating a prominent role for O antigen in GVIN1 coatomer formation and also explaining why the Δ*wbpm* mutant strain is resistant to GBP1-independent, i.e. GVIN1-dependent, host defense (Fig. 3E). In contrast to GVIN1 coatomers the frequency of GBP1 coatomer formation was comparable between WT and Δ*wbpm B. thailandensis* mutant bacteria (Fig. S8C), suggesting that the underlying mechanisms for GBP1 and GVIN1 coatomer formation are distinct from each other.

Ectopically expressed GVIN1 formed coatomers around *B. thailandensis* both in unprimed and IFNγ-primed cells (Fig. 5F). However, a reduction in *B. thailandensis* actin tail formation required IFNγ priming in addition to ectopic GVIN1 expression both in HeLa and T24 cells (Fig. 5C-D and 5G-H), indicating that GVIN1 requires an IFNγ-inducible co-factor for host defense. Because ectopically expressed GVIN1 restricted bacterial actin tail formation in IFNγ-primed GBP1^KO^ HeLa as well as GVIN1/GBP1^DKO^ T24 cells (Fig. 5D and 5H), GBP1 can be excluded as an essential co-factor for GVIN1 function.

### The *Burkholderia* actin nucleator BimA is displaced from the bacterial pole by both GBP1- and GVIN1-dependent host defense pathways

We previously demonstrated that GBP1 coatomers on the surface of *S. flexneri* in HeLa cells are sufficient to disrupt the polar localization of the *Shigella* actin nucleator IcsA (*13*). In contrast to IcsA, we found that the polar localization of the *Burkholderia* actin nucleator BimA, tagged with the DYKDDDDK peptide (FLAG), remained unchanged in GBP1 coated bacteria residing within HeLa cells (Fig. S9A-B). These findings were expected considering that GBP1 coatomers are similarly insufficient to block *B. thailandensis* actin tail formation in Hela cells (Fig. 1C). Next, we monitored BimA localization in T24 cells, which in contrast to HeLa cells dramatically reduce actin tail formation by *B. thailandensis* upon IFNγ priming (Fig. 1F). We observed that the number of bacteria with detectable BimA was dramatically diminished in IFNγ-primed WT T24 cells (Fig. 6A-B). Because ‘actin clouds’ associated with dispersed BimA expression were not detectable on GVIN1-coated *B. thailandensis* inside IFNγ-primed cells (Fig. 5F), the loss of BimA detection likely reflects a loss of BimA surface expression rather than mere BimA displacement from the bacterial pole.

**Fig. 6.**
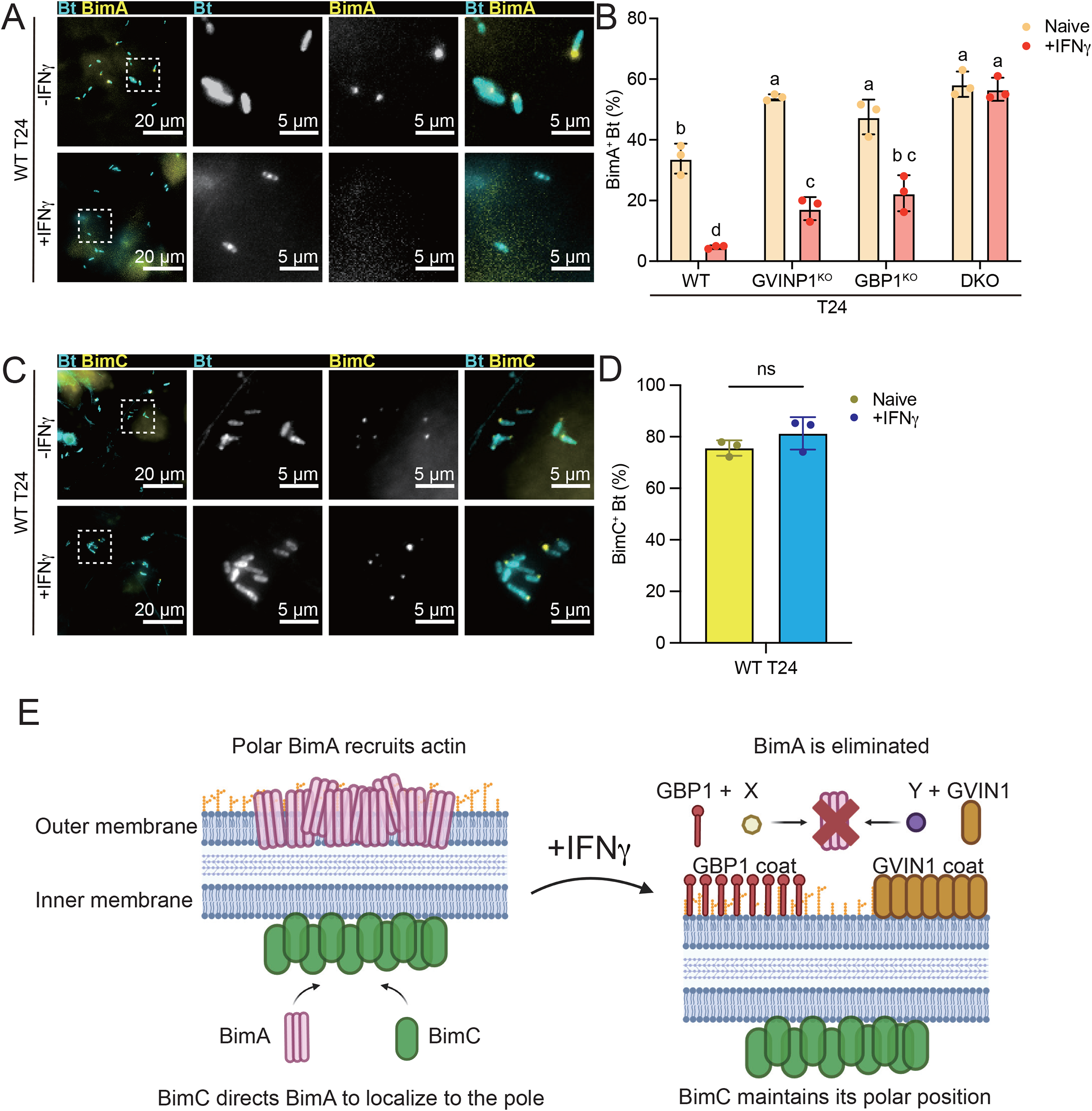
BimA is displaced from the bacterial pole via both GBP1- and GVIN1-dependent pathways, whereas BimC remains unaffected. (A-B) Naïve or 200 U/mL IFNγ primed WT, GBP1^KO^, GVIN^KO^, or GBP1^KO^ GVIN^KO^ DKO T24 cells were infected with *B. thailandensis* expressing FLAG-tagged BimA at an MOI of 100. Cells were fixed at 8 hours post infection and stained for FLAG-BimA. Percentages of BimA positive *B. thailandensis* were quantified (B). (C-D) Naïve or 200 U/mL IFNγ primed WT T24 cells were infected with *B. thailandensis* expressing GFP-tagged BimC at an MOI of 100. Cells were fixed at 8 hours post infection. Percentages of GFP-BimC positive *B. thailandensis* were quantified (D). All bar graphs show 3 independent biological replicates, presented as mean ± SD. Two-way ANOVA with Tukey’s multiple comparison tests were performed for (B), and unpaired t-test was performed for (D). Specific p-values indicated as follows: (B) columns not sharing any letters means significantly different (p<0.05); (D) ns, not significant. (E) Schematic depiction of the model, Polar BimA that recruits actin is eliminated by the GBP1 pathway and GVIN1 pathway together in IFNγ primed T24 cells.

In contrast to the outer membrane protein BimA, the BimA-interacting inner membrane protein BimC (*33*) retained its polar localization in IFNγ-primed WT T24 cells (Fig. 6C-D). Polar BimA localization was partially restored in IFNγ-primed GBP1^KO^ and GVIN1^KO^ T24 cells (Fig. 6B). In GVIN1/GBP1^DKO^ the percentage of BimA-positive bacteria was identical between unprimed and IFNγ-primed cells (Fig. 6B). Together, these observations demonstrated that both the GBP1 and GVIN1 host defense pathways eliminate BimA from the bacterial outer membrane to block actin tail formation (Fig. 6E).

## DISCUSSION

IFN-inducible GTPases are potent executioners of cell-autonomous immune programs targeting viral, bacterial, fungal, and protozoan intracellular pathogens. Previous work focused on the function of three of these GTPases families, the Mx proteins, the IRGs, and the GBPs (*17–20, 34, 35*). While *GVIN* genes encoding members of the fourth family of IFN-inducible GTPases are present in most vertebrate genomes and have previously been shown to be expressed in mice (*36*), their biological function has remained enigmatic. Our current work demonstrates that human GVIN1, the sole human member of the GVIN family, forms a coatomer around *B. thailandensis* and interferes with bacterial actin tail formation. Our observations thus establish GVIN GTPases as a new class of host defense proteins.

Previous work demonstrated that human GBP1 can form coatomers on the surface of numerous Gram-negative bacteria to combat intracellular infections (*13, 30–32*). Structure-function analyses of GBP1 demonstrated that its coatomer-forming activity can be uncoupled from its contribution to noncanonical inflammasome activation (*37*). Instead of playing a role in cell death, our previous work supports a model in which the GBP1 coatomer acts like an emulsifier and disrupts the chemo-mechanical barrier on the bacterial surface established by the polysaccharide O antigen portion of bacterial LPS (*13*). In concert with additional antimicrobial molecules the emulsification of the LPS barrier by GBP1 can promote the killing of encased bacteria (*13, 38*). On its own the GBP1 coatomer appears to be neither bacteriostatic nor bactericidal (*21*), yet its detergent-like activity is sufficient to increase the diffusion rate of the *Shigella* outer membrane protein IcsA, thereby interfering with IcsA’s critical role in nucleating actin at one bacterial pole and establishing actin-based motility (*13*). In contrast to the *Shigella* IcsA protein, we show here that the functionally related BimA protein of *B thailandensis* is impervious to the disruption of the O antigen layer. GBP1 must therefore block *B. thailandensis* actin tail formation through a mechanism distinct from the one it employs against *Shigella*. Consistent with this model, our data indicate that one or more as-yet-unidentified GBP1 co-factors are required for GBP1 coatomers to block actin tail formation specifically by *B thailandensis*.

Like GBP1 coatomers, GVIN1 coatomers require additional co-factors to effectively block *B. thailandensis* actin formation. The nature of these co-factors will need to be determined in future studies. However, because ectopic expression of GVIN1 but not GBP1 in HeLa cells reduces *B. thailandensis* actin tail formation, we can already conclude that HeLa cells express essential GVIN1 but not GBP1 co-factors. Accordingly, we can infer that GBP1 and GVIN1 use distinct sets of co-factors. Despite these clear differences, both GBP1 and GVIN1 coatomers, in cooperation with their respective co-factors, function similarly by removing BimA from the bacterial outer membrane to disrupt actin tail formation by *B. thailandensis.* The polar localization of the inner membrane protein BimC, on the other hand is undisturbed, suggesting that both GBP1 and GVIN1 coatomers specifically disrupt modalities of the outer bacterial membrane.

The basic principles underlying the assembly of GBP1 coatomers have been established. Upon GTP binding and hydrolysis GBP1 monomers undergo conformational changes and form dimers (*39, 40*). These dimers can polymerize (*41*), a process that is aided by the presence of bacterial LPS (*13, 37*). The polymers dock to the surface of Gram-negative bacteria, penetrate the bacterial polysaccharide barrier, and release thousands of GBP1 dimers into the bacterial outer membrane to form a GBP1 coat on the bacterial surface (*13, 18, 31*). Although our structural modelling suggests that GVIN1 GTPases may also form dimers, whether GVIN1 coatomers assemble through a multi-step process that is analogous to GBP1 coatomer formation will require future investigations.

Our studies did not systematically explore whether GVIN1 targets intracellular pathogens other than *B. thailandensis,* although this seems likely. Remarkably, we found that O antigen deficient *B. thailandensis* rough mutants had become resistant to GVIN1-mediated host defense, which suggests that GVIN1 either detects moieties present in *B. thailandensis* O antigen or that O antigen helps to stabilize the GVIN1 coat. This observation is also noteworthy in the context of reports showing that clinically important *Burkholderia* species such as *Burkholderia cenocepacia* and *Burkholderia multivorans* isolated from chronically infected cystic fibrosis patients tended to acquire mutations that alter or remove O antigen (*42–44*). While speculative, changes in O antigen composition and expression may provide these clinical *Burkholderia* isolates with a formidable fitness advantage by obtaining resistance to GVIN1.

ISGs play crucial roles in executing cell-autonomous immune responses. However, only a limited number of these ISGs have been thoroughly characterized, and there is little understanding of how they functionally interact with one another. Here, we characterize GVIN proteins as novel executioners of cell-autonomous immunity. Like related GBP1 proteins, human GVIN1 proteins encase their bacterial target, spotlighting coatomer formation as a fundamental mechanism of antibacterial cell-autonomous defense. Furthermore, we show that both GBP1 and GVIN1 coatomers collaborate with other ISGs, highlighting ISG synergy as a principal operating mechanism of antimicrobial coatomer activity. Elucidating the formation and defensive role of GVIN1 coatomers, along with potential microbial evasion strategies, could offer valuable insights into host-pathogen interactions and aid in developing novel therapies for infectious diseases.

## MATERIALS AND METHODS

### Cell lines and culture

HeLa (ATCC #CCL-2), A549 (ATCC #CCL-185), and 293T (ATCC #CRL-3216) cell lines were cultivated in Dulbecco’s Modified Eagle Medium (DMEM, Gibco) supplemented with 10% heat inactivated fetal bovine serum (Omega Scientific), 1% MEM Non-Essential Amino Acids Solution (MEM NEAA, Gibco), and 55 µM 2-Mercaptoethanol (Gibco). T24 (ATCC #HTB-4) cells were cultivated in McCoy’s 5A (Modified) Medium (Gibco) supplemented with 10% heat inactivated fetal bovine serum. THP-1 (ATCC #TIB-202) and U-937 (ATCC #CRL-1593.2) cells were cultivated in Roswell Park Memorial Institute 1640 (RPMI 1640, Gibco) supplemented with 10% heat inactivated fetal bovine serum, and 55 µM 2-Mercaptoethanol (Gibco). THP-1 and U-937 monocytes were differentiated to macrophages by incubation with 50ng/mL phorbol 12-myristate 13-acetate (PMA, Sigma-Aldrich) for 48 hours. All cell lines were grown at 37 °C with 5% CO_2_. All cell lines were authenticated by the Duke University DNA Analysis Facility using GenePrint10 (Promega) and were routinely tested for mycoplasma.

### Knock out cell lines

The knockout (KO) GBP1, GVIN1, and GBP6 T24 cell lines were generated by the Duke Functional Genomics core using CRISPR/Cas9 technology. CRISPR sgRNAs targeting GBP1, GVIN1, and GBP6 were designed using ChopChop and ordered as modified synthetic sgRNAs from Synthego (*45*). For GBP1, the sgRNA sequence 5’-TGCCATTACACAGCCTATGG-3’ was used. For GVIN1, the sgRNA sequence 5’-TGTACATTGTGGCTGTAGCG-3’ was used. For GBP6 the sgRNA sequences 5’-TCTGGCAGTGACTTATGTAG-3’ was used. 5.6 x 10^4^ T24 cells were electroporated with 6 pmol TrueCut Cas9 protein v2 (ThermoFisher Scientific) complexed with 36 pmol sgRNA using the Neon system (ThermoFisher Scientific) with the following settings: 1600 V, 10 ms, 3 pulses. Cells were recovered and expanded, followed by PCR sequencing and analysis using Inference of CRISPR Edits (ICE) to determine KO efficiencies for each individual sgRNA (*46*). For each KO cell line, single clones were isolated by plating at limited dilution in 96 well plates. Clones were expanded for approximately 10-14 days before screening by PCR-sequencing to confirm that the edit resulted in a frame-shift mutation. The GBP1^KO^ HeLa cell line was described previously (*13, 47*). The CASP4^KO^ THP-1 cell line (*48*) was a gift from Dr. Veit Hornung, University of Munich, Germany, and Dr. Eva Frickel, University of Birmingham, United Kingdom.

### Bacterial strains

Wild-type *B. thailandensis* strain E264 carrying a GFP expression construct was a gift from Dr. Edward Miao, Duke University, USA. For the BimA studies, a *B. thailandensis* strain E264 was used with BimA chromosomally replaced with Flag-BimA (*10*), which was a gift from Dr. Matthew D. Welch, University of California, Berkeley, USA. For the BimC studies, wild-type *B. thailandensis* strain E264 was transformed with pME6032-GFP-BimC (*33*), a gift from Dr. Feng Shao, National Institute of Biological Sciences, Beijing, China. The *wbpM* (BTH_I1483) gene was deleted from the native locus of *B. thailandensis* by allelic exchange using homologous recombination. Regions (∼500 bp) flanking *wbpM* with a chloramphenicol resistance cassette (Cm^r^) in the middle were cloned into a pEXKm5 suicide vector (*49*), which was kindly provided by Dr. Peggy Cotter, University of North Carolina, Chapel Hill, USA. The recombinant plasmid was transformed into *B. thailandensis*. Briefly, an overnight culture of *B.thailandensis* was back-diluted and grown in minimal media, M63 for 4 hours (M63 5X salt solution was made by dissolving 15 g KH_2_PO_4_, 35 g KHPO_4_, 10 g (NH_4_)_2_SO_4_, and 2.5 mL of a 16 nM FeSO_4_ in 1 L of water and the pH was adjusted to 7.0; to make 1 L of the final M63 medium, 10 mL of 1% casamino acids was added to 780 mL water and autoclaved. Sterile 200 mL of the M63 5X salt solution, 10 mL of 20% glucose, 1 mL of 1 M MgSO_4_, and 8 mL of 50% glycerol was added). *B. thailandensis* mixed with linearized pEXKm5 suicide vector containing *wbpM* flanking sequences were grown overnight in minimal media. The next day, the culture was pelleted and spread on agar plates to select for chloramphenicol resistance. Double crossover was confirmed by PCR using Cm^r^-F (5’-TCAGTCAGCGGCCGCCACGAGGCCCTTTCGTCTTCGAATAA-3’) and Cm^r^-R (5’-AGTCACTGCGGCCGCTTGCGCTCACTGCCCGCTTTCCAG-3’) as well as wbpM-Flnk-F (5’-TCGATGACCGTCGCGGATTGC-3’) and wbpM-Flnk-R (5’ATGGTCGCGTTCATGATGTTCGC3’) primer sets. *S. flexneri ΔipaH9.8* strain carrying pGFPmut2 was previously reported (*21*). Bacterial cultures were supplemented with the following antibiotics when required: carbenicillin (50 μg/mL), kanamycin (50 μg/mL), chloramphenicol (25 μg/mL), gentamicin (25 μg/mL). Antibiotic concentrations were maintained throughout infection experiments.

### LPS visualization

Overnight grown bacterial cultures were lysed with RIPA buffer (Thermo Fisher Scientific,89900) and checked for protein concentration using bicinchoninic acid (BCA) assay kit (Thermo Fisher Scientific). Aliquots of 20 μg samples were diluted in Laemmli buffer and applied to a 4–20% Mini-PROTEAN® TGX™ Precast Protein Gels (Biorad, 4561094). The gel was processed with the Pro-Q Emerald 300 LPS stain kit (Thermo Fisher Scientific, P20495), according to the manufacturer’s protocol. The LPS bands were visualized under UV light using the Azure 500 imaging system (Azure Biosystems).

### Bacterial infection

Cell lines (T24, HeLa, A549, THP-1, and U-937) were seeded in 24-well plate (immunofluorescence and CFU assays) or 96-well plates (LDH assay) and treated with 200 U/ml IFNγ (Sigma-Aldrich) overnight before infection. For *B. thailandensis* infections, all *B. thailandensis* strains were grown at 37°C on lysogeny broth (LB) agar plates. 5 mL of LB supplemented with appropriate antibiotics was inoculated with one *B. thailandensis* colony and incubated overnight at 37°C with shaking. On the day of infection, the overnight culture was diluted 1:50 in 5 mL of fresh LB with antibiotics and grown at 37°C with shaking for 3–4 hours until the optical density at 600 nm (OD600) reached 0.6–0.8. The bacterial cultures were pelleted by centrifugation at 5000 rpm for 3 minutes at room temperature (RT), resuspended in 1 mL of phosphate-buffered saline (PBS, Gibco), and centrifuged again under the same conditions. The washed bacteria were then resuspended in cell culture media containing antibiotics and further diluted in cell culture media to the desired concentration. T24, HeLa and A549 cells were infected at a multiplicity of infection (MOI) of 100, THP-1 and U-937 cells were infected at a MOI of 20. Following centrifugation at 700 × g for 10 minutes at RT, infected cells were incubated at 37°C with 5% CO_2._ At 4 hpi, cells were washed twice with Hank’s Balanced Salt Solution (HBSS, Gibco) to remove extracellular bacteria. Cells were then incubated in media containing chloramphenicol for 2 hours to eliminate remaining extracellular bacteria. Afterwards, the culture media was replaced with antibiotic-free media. The infection was terminated at 8 hpi for T24, HeLa, and A549 cells, and at 6 hpi for THP-1 and U-937 cells. For *S. flexneri* infections, bacteria were grown at 37°C on tryptic soy broth (TSB) agar plates containing 0.01% Congo red. 5 mL of TSB supplemented with appropriate antibiotics was inoculated with one *S. flexneri* colony and incubated overnight at 37°C with shaking. On the day of infection, the overnight culture was diluted 1:50 in 5 mL of fresh TSB with antibiotics and grown at 37°C with shaking for about 2 hours until the OD600 reached 0.5-0.7. The bacterial cultures were pelleted by centrifugation at 5000 rpm for 3 minutes RT, resuspended in the 990 µL PBS supplement with 10 µL of 0.1 mg/mL polyD-lysine (Sigma-Aldrich) solution, and then incubated at 37°C with shaking for 15 minutes. The *S. flexneri* were pelleted again by centrifugation at 5000 rpm for 3 minutes RT. The washed bacteria were then resuspended in cell culture media containing antibiotics and further diluted in cell culture media to the desired concentration. T24, HeLa cells were infected at an MOI of 10, followed by centrifugation at 700 × g for 10 minutes at RT. Infected cells were incubated at 37°C with 5% CO_2._ 30 minutes post-infection, cells were washed twice with HBSS to remove extracellular bacteria. Cells were then incubated in culture media containing gentamicin to eliminate extracellular bacteria. The infection was terminated at 2 hours and 30 minutes post infection for T24 and HeLa cells.

For CFU and plaque assay T24 cells and KO cell lines were infected with *B. thailandensis* at an MOI of 10 and 0.1, respectively. At 4h hpi, cells were washed with 1X PBS and incubated with media containing 500*_μ_*g/mL gentamicin for 1h. Cells were then washed thoroughly with 1X PBS and cultured with fresh media containing 20*_μ_*g/mL gentamicin for the remainder of the experiment. Cells were lysed at 8h hpi and lysates were plated onto LB agar plates too enumerate CFUs. For plaque assays cells were washed at at 18 hpi and immediately fixed and stained with 1% crystal violet solution (Sigma) for 15 min.

### Expression constructs and lentivirus production

An anhydrotetracycline-inducible GVIN1 plasmid was constructed using Gateway assembly (Thermo Fisher) into the pMVP cloning system. The pMVP Cloning System was a gift from Christopher Newgard (Addgene kit #1000000155) (*50*). The third ORF of GVIN1 sequence without stop codon flanked by attB4 and attB3r recombination sites was amplified via PCR using primers, CloneGVINP1F3 (5’-GGGGACAACTTTTCTATACAAAGTTGCCACCATGCAGTGGATCTTCCAGTC-3’), and CloneGVINP1nostopR1 (5’-GGGGACAACTTTATTATACAAAGTTGTCTGTTTTTTCAAGTCTTCAAGCAC-3’) with genomic DNA from T24 cells as the template. The PCR product was cloned into the pDONR221-P4r-P3r vector (NovoPro) using BP Clonase™ II (Thermo Fisher), generating a GVIN1 entry clone. Subsequently, the following entry clones were assembled in an LR recombination reaction: pDONR221-R4-R3-GVIN1, Tre promoter entry clone (Addgene Plasmid #121691), HA tag-polyA + CMV-TETa-polyA entry clone (Addgene Plasmid #121802).Entry clone plasmids were added with the pMVP/Lenti/Neo-DEST vector (Addgene Plasmid #121850), and LR Clonase™ II Plus enzyme mix (Thermo Fisher) together to generate the final tetracycline-inducible HA-tagged GVIN1 expression lentiviral construct (pMVP-lenti-neo-largeORFGVIN1-HA). Lentiviral particles were produced by transfecting lentivirus plasmids with packaging vectors into HEK293T cells using the TransIT™ 293 transfection reagent (Mirus Bio) according to the manufacturer’s instructions. Each well of a 6-well plate containing 293T cells was transfected with 1 µg of plasmid, 750 ng of pSPAX2, and 250 ng of VSVG. At 24 hours post-transfection, the media were replaced with DMEM. The supernatants containing lentiviral particles were collected at 48- and 72-hours post-transfection. Supernatants were pooled and filtered through 0.45-µm nylon filters (Corning). For lentiviral transduction, 293T cells were diluted to a concentration of 3.33 × 10⁴ cells/mL in media containing 10 µg/mL polybrene. A total of 500 µL of lentivirus was added to each well of a 6-well plate containing recipient cells, followed by the addition of 1.5 mL of diluted cells. The plates were incubated at 37°C with 5% CO₂ for 48 hours. Post transduction, cells were selected in media containing geneticin (Gibco, 500 μg /mL for HeLa, 1000 μg /mL for T24). An anhydrotetracycline-inducible GBP1 plasmid was described previously (*13, 37, 47*). GBP1^KO^ HeLa cells and GBP1^KO^ T24 cells were stably transduced with this anhydrotetracycline-inducible mCherry-hGBP1 expression system, as described above.

### Immunofluorescence microscopy

Post-infection, cells were fixed in 4% paraformaldehyde in PBS at RT for 20 minutes. Fixed cells were permeabilized with 0.25% Triton X-100 in PBS for 10 minutes at RT. Permeabilized cells were then blocked with blocking buffer (5% bovine serum albumin and 0.3 M glycine in PBS) at RT for 45 minutes. Samples were incubated with primary antibodies diluted in blocking buffer at RT for 1 hour. Primary antibodies were diluted in blocking buffer as follows: rabbit recombinant monoclonal anti-GBP1 (1:150; Abcam, ab131255); mouse monoclonal ANTI-FLAG® M2 (1:1000; Sigma-Aldrich, F1804); mouse monoclonal anti-HA tag [HA.C5] (1:500; Abcam, ab18181); recombinant rabbit monoclonal Anti-LPS [CK3 (BP1 7F7)] (absolute antibody, Ab03071-23.0). Following primary antibody incubation, samples were washed three times in 0.05% Triton X-100 in PBS for 5 minutes at RT. Subsequently, samples were incubated with secondary antibodies diluted in blocking buffer at RT for 1 hour. Secondary antibodies and phalloidin were prepared in blocking buffer as follows: Alexa Fluor™ 568-conjugated donkey anti-rabbit IgG (H+L; 1:1000; Thermo Fisher, A10042), Alexa Fluor™ 405-conjugated goat anti-mouse IgG (H+L; 1:1000; Thermo Fisher, A31553), Alexa Fluor™ 568-conjugated goat anti-mouse IgG (H+L; 1:1000; Thermo Fisher, A-11004), Alexa Fluor™ 488-conjugated goat anti-rabbit IgG (H+L; 1:1000; Thermo Fisher, A-11008), Alexa Fluor™ 660 phalloidin (1:400; Thermo Fisher, A22285) for F-actin visualization. Cells were washed four times in 0.05% Triton X-100 in PBS for 5 minutes at RT. Coverslips with cells were mounted on glass slides using a mixture of Mowiol 4-88 and para-phenylenediamine antifading agent (9:1 ratio) to preserve fluorescence. Images were acquired using a Zeiss Axio Imager microscope equipped with a 63×/1.4 NA oil immersion lens. For each condition, at least six fields of view were captured to analyze at least 100 bacteria. Z-stack images were taken at 1 µm intervals. Bacteria with protein signals surrounding at least 50% of its visible surface were counted as targeted.

### Western blotting

For cell lysis and protein extraction, cells were washed twice with Hank’s Balanced Salt Solution (HBSS) and lysed in RIPA buffer (Sigma-Aldrich) supplemented with 4 U/ml DNase I (New England Biolabs) and protease inhibitor cocktail (Sigma-Aldrich). Lysates were incubated at 4°C for 30 minutes, followed by centrifugation at 20,000 × g for 10 min at 4°C. Protein concentrations were quantified using a bicinchoninic acid (BCA) assay kit (Thermo Fisher Scientific) and normalized prior to analysis. Normalized samples were mixed with Laemmli buffer containing 5% β-mercaptoethanol and denatured by heating at 95°C for 10 min. Protein samples were resolved on 4–20% Mini-PROTEAN TGX Stain-Free precast gels (Bio-Rad) using Tris-glycine running buffer. Proteins were transferred to PVDF membranes using the Tank Transfer System or Trans-Blot Turbo Transfer System (both Bio-Rad). Primary antibodies diluted in blocking solution (5% non-fat dry milk in 0.1% TBS-T) were incubated with membranes overnight at 4°C. After three washes with TBS-T (0.1% Tween-20 detergent in tris-buffered saline), membranes were incubated with species-matched horseradish peroxidase (HRP)-conjugated secondary antibodies for 1 hour at room temperature. After four washes with TBS-T, protein bands were visualized using Amersham ECL Prime Western Blotting Detection Reagent (Cytiva) and imaged on an Azure 500 imaging system (Azure Biosystems).

### LDH assay

Cell death was evaluated using the lactate dehydrogenase (LDH) release assay (CytoTox-ONE™ Homogeneous Membrane Integrity Assay (Promega)) to measure cell membrane damage. Briefly, an equal volume (50-100 μL) of CytoTox-ONE™ reagent was added to each well in a 96 well plate with cell samples post-infection. The reaction mixture was incubated in the dark at RT for 10 min to allow enzymatic conversion of resazurin to resorufin by released lactate dehydrogenase (LDH). Fluorescence intensity was measured using a plate reader (Biotek Synergy H1, Agilent).

### RNA sequencing

T24, HeLa, A549 cells were each plated in two wells of a 6 well plate (Corning) and allowed to grow overnight in standard media and culture conditions (described above). After 24 hours, media was replaced with fresh standard media (control unprimed well) or media containing 100 U/ml IFNγ (primed well). Six hours after priming, RNA was harvested for analysis. THP1 and U937 cells were differentiated into macrophages by treating with 50 ng/ul PMA in standard media for 48 hours and then cultured for another 48 hours in standard media without PMA. Then, media was replaced with fresh standard media (control unprimed well) or media containing 100 U/ml IFNγ (primed well). Six hours after priming, RNA was harvested for analysis. RNA was isolated using RNeasy Mini Kit (Qiagen) following manufacturer’s instructions. The RNA library was prepared using KAPA mRNA Hyperprep and sequenced on the Illumina NovaSeq 6000. RNA-seq data was processed using the fastp toolkit (*51*) to trim low-quality bases and sequencing adapters from the 3’ end of reads, then mapped to GRCh38 (downloaded from Ensembl, version 106) (*52*) using the STAR RNA-seq alignment tool (*53*) and reads aligning to a single genomic location were summarized across genes. For genes having an overlap of at least 10 reads, gene counts were normalized, and differential expression was carried out using the DESeq2 (*54*) and Bioconductor (*55*) package implemented for the R programming environment. Consistent with the recommendation of the DESeq authors, independent filtering (*56*) was utilized prior to calculating adjusted p-values (*57*) and moderated log2 fold-changes were derived using the ashr package (*58*).

### siRNA knockdown

Gene knockdown in T24 cell lines was achieved by transfecting cells with siGENOME siRNA Reagents (Horizon) using Lipofectamine® RNAiMAX Reagent (Thermo Fisher Scientific) following the manufacturer’s instructions. In brief, T24 cells were seeded in 24-well plates to achieve 60–80% confluency at the time of transfection. siRNA-lipid complexes were prepared in Opti-MEM® Reduced Serum Medium (Gibco). The siRNA and lipofectamine reagents were diluted separately in Opti-MEM® medium before mixing the dilutions in a 1:1 volume ratio. Following incubation at room temperature for 5 minutes to allow formation of the siRNA-lipid complexes, the reaction mix was added to the T24 cells. The final transfection mixture for each well contained 5 pmol siRNA and 1.5 µL Lipofectamine in 50 µL siRNA-lipid Opti-MEM® complex. After 48 h, transfected cells were used for the infection screening. The siRNA sequences used are listed in table 1 of our supplementary data.

### Protein structure prediction and modelling

Structural models of GVIN1 were predicted with the Alphafold server (https://alphafoldserver.com) using the Alphafold3 algorithm (*27*). Therefore, the protein sequence was provided as input, and for the dimer model, guanosine triphosphate and divalent magnesium cations were chosen as ligands and ions, respectively. To define the domain structure, structural alignments were done with Foldseek (*28*). Structural analysis and comparison were conducted using the PyMOL molecular graphics system (version 3.1.3.1, Schrödinger). PAE plots were visulalized with UCSF ChimeraX (version 1.9).

## Statistical analysis

All graphical representations and statistical analyses were performed using GraphPad Prism 10. All experimental data were derived from at least 3 independent experiments and are presented as mean ± standard deviation (SD) in bar/line graphs. Parametric statistical comparisons were conducted using Two-tailed unpaired Student’s t-test for pairwise comparisons and Two-way ANOVA with Tukey’s multiple comparisons test for factorial experimental designs. p-values are indicated as follows: *p < 0.05, **p < 0.01, ***p < 0.001, ****p < 0.0001.

## Data storage

The RNA-seq data discussed in this publication have been deposited in NCBI’s Gene Expression Omnibus (*59*) and will be accessible through GEO Series accession number GSE288527 (https://www.ncbi.nlm.nih.gov/geo/query/acc.cgi?acc=GSE288527). The numerical values of all other quantified data are deposited as source data.

## ACKNOWLEDGEMENTS

This work was supported by National Institutes of Health grants AI139425 (to JC). The funders had no role in study design, data collection and interpretation, or the decision to submit the work for publication. We would like to thank members of the Coers lab as well as the labs of Drs. Clare Smith, David Tobin, and Edward Miao for providing valuable feedback. We thank Dr. Peggy Cotter at the University of Chapel Hill, North Carolina and Dr. Matt Walch at the University of California, Berkeley for providing reagents and invaluable advice on all things related to *Burkholderia*. We thank the Duke University School of Medicine for the use of the Functional Genomics and Sequencing and Genomic Technologies Shared Resource, which provided gene editing and RNA sequencing services.

## REFERENCES

1. V. K. Koseoglu, H. Agaisse, Evolutionary Perspectives on the Moonlighting Functions of Bacterial Factors That Support Actin-Based Motility. mBio 10, (2019).

2. R. L. Lamason, M. D. Welch, Actin-based motility and cell-to-cell spread of bacterial pathogens. Curr Opin Microbiol 35, 48–57 (2017).

3. M. L. Bernardini, J. Mounier, H. d’Hauteville, M. Coquis-Rondon, P. J. Sansonetti, Identification of icsA, a plasmid locus of Shigella flexneri that governs bacterial intra-and intercellular spread through interaction with F-actin. Proc Natl Acad Sci U S A 86, 3867–3871. (1989).

4. M. B. Goldberg, O. Barzu, C. Parsot, P. J. Sansonetti, Unipolar localization and ATPase activity of IcsA, a Shigella flexneri protein involved in intracellular movement. Infect Agents Dis 2, 210–211 (1993).

5. C. Egile et al., Activation of the CDC42 effector N-WASP by the Shigella flexneri IcsA protein promotes actin nucleation by Arp2/3 complex and bacterial actin-based motility. J Cell Biol 146, 1319–1332 (1999).

6. R. P. Mauricio, C. M. Jeffries, D. I. Svergun, J. E. Deane, The Shigella Virulence Factor IcsA Relieves N-WASP Autoinhibition by Displacing the Verprolin Homology/Cofilin/Acidic (VCA) Domain. J Biol Chem 292, 134–145 (2017).

7. T. Suzuki, H. Miki, T. Takenawa, C. Sasakawa, Neural Wiskott-Aldrich syndrome protein is implicated in the actin-based motility of Shigella flexneri. EMBO J 17, 2767–2776 (1998).

8. C. Sitthidet et al., Actin-based motility of Burkholderia thailandensis requires a central acidic domain of BimA that recruits and activates the cellular Arp2/3 complex. J Bacteriol 192, 5249–5252 (2010).

9. J. M. Stevens et al., Actin-binding proteins from Burkholderia mallei and Burkholderia thailandensis can functionally compensate for the actin-based motility defect of a Burkholderia pseudomallei bimA mutant. J Bacteriol 187, 7857–7862 (2005).

10. E. L. Benanti, C. M. Nguyen, M. D. Welch, Virulent Burkholderia species mimic host actin polymerases to drive actin-based motility. Cell 161, 348–360 (2015).

11. N. P. Giordano, M. B. Cian, Z. D. Dalebroux, Outer Membrane Lipid Secretion and the Innate Immune Response to Gram-Negative Bacteria. Infect Immun 88, (2020).

12. E. R. Rojas et al., The outer membrane is an essential load-bearing element in Gram-negative bacteria. Nature 559, 617–621 (2018).

13. M. Kutsch et al., Direct binding of polymeric GBP1 to LPS disrupts bacterial cell envelope functions. EMBO J 39, e104926 (2020).

14. R. C. Sandlin et al., Avirulence of rough mutants of Shigella flexneri: requirement of O antigen for correct unipolar localization of IcsA in the bacterial outer membrane. Infect Immun 63, 229–237 (1995).

15. R. C. Sandlin, M. B. Goldberg, A. T. Maurelli, Effect of O side-chain length and composition on the virulence of Shigella flexneri 2a. Mol Microbiol 22, 63–73 (1996).

16. J. L. Casanova, J. D. MacMicking, C. F. Nathan, Interferon-gamma and infectious diseases: Lessons and prospects. Science 384, eadl2016 (2024).

17. D. Pilla-Moffett, M. F. Barber, G. A. Taylor, J. Coers, Interferon-inducible GTPases in host resistance, inflammation and disease. Journal of molecular biology, (2016).

18. M. Kutsch, J. Coers, Human guanylate binding proteins: nanomachines orchestrating host defense. FEBS J, (2020).

19. M. Kirkby, D. Enosi Tuipulotu, S. Feng, J. Lo Pilato, S. M. Man, Guanylate-binding proteins: mechanisms of pattern recognition and antimicrobial functions. Trends Biochem Sci 48, 883–893 (2023).

20. Y. Rivera-Cuevas, B. Clough, E. M. Frickel, Human guanylate-binding proteins in intracellular pathogen detection, destruction, and host cell death induction. Curr Opin Immunol 84, 102373 (2023).

21. A. S. Piro et al., Detection of Cytosolic Shigella flexneri via a C-Terminal Triple-Arginine Motif of GBP1 Inhibits Actin-Based Motility. mBio 8, (2017).

22. M. P. Wandel et al., GBPs Inhibit Motility of Shigella flexneri but Are Targeted for Degradation by the Bacterial Ubiquitin Ligase IpaH9.8. Cell Host Microbe 22, 507–518 e505 (2017).

23. M. Dilucca, S. Ramos, K. Shkarina, J. C. Santos, P. Broz, Guanylate-Binding Protein-Dependent Noncanonical Inflammasome Activation Prevents Burkholderia thailandensis-Induced Multinucleated Giant Cell Formation. mBio 12, e0205421 (2021).

24. D. E. Place et al., Interferon inducible GBPs restrict Burkholderia thailandensis motility induced cell-cell fusion. PLoS Pathog 16, e1008364 (2020).

25. C. T. French et al., Dissection of the Burkholderia intracellular life cycle using a photothermal nanoblade. Proc Natl Acad Sci U S A 108, 12095–12100 (2011).

26. J. Jumper et al., Highly accurate protein structure prediction with AlphaFold. Nature 596, 583–589 (2021).

27. J. Abramson et al., Accurate structure prediction of biomolecular interactions with AlphaFold 3. Nature 630, 493–500 (2024).

28. M. van Kempen et al., Fast and accurate protein structure search with Foldseek. Nature biotechnology 42, 243–246 (2024).

29. O. Daumke, G. J. Praefcke, Invited review: Mechanisms of GTP hydrolysis and conformational transitions in the dynamin superfamily. Biopolymers 105, 580–593 (2016).

30. S. Zhu et al., Native architecture of a human GBP1 defense complex for cell-autonomous immunity to infection. Science 383, eabm9903 (2024).

31. T. Kuhm et al., Structural basis of antimicrobial membrane coat assembly by human GBP1. Nat Struct Mol Biol 32, 172–184 (2025).

32. M. Weismehl et al., Structural insights into the activation mechanism of antimicrobial GBP1. EMBO J 43, 615–636 (2024).

33. Q. Lu, Y. Xu, Q. Yao, M. Niu, F. Shao, A polar-localized iron-binding protein determines the polar targeting of Burkholderia BimA autotransporter and actin tail formation. Cell Microbiol 17, 408–424 (2015).

34. J. Dockterman, J. Coers, How did we get here? Insights into mechanisms of immunity-related GTPase targeting to intracellular pathogens. Curr Opin Microbiol 69, 102189 (2022).

35. O. Haller, H. Arnheiter, J. Pavlovic, P. Staeheli, The Discovery of the Antiviral Resistance Gene Mx: A Story of Great Ideas, Great Failures, and Some Success. Annu Rev Virol 5, 33–51 (2018).

36. T. Klamp, U. Boehm, D. Schenk, K. Pfeffer, J. C. Howard, A giant GTPase, very large inducible GTPase-1, is inducible by IFNs. J Immunol 171, 1255–1265 (2003).

37. M. S. Dickinson et al., LPS-aggregating proteins GBP1 and GBP2 are each sufficient to enhance caspase-4 activation both in cellulo and in vitro. Proc Natl Acad Sci U S A 120, e2216028120 (2023).

38. R. G. Gaudet et al., A human apolipoprotein L with detergent-like activity kills intracellular pathogens. Science 373, (2021).

39. M. Wehner, S. Kunzelmann, C. Herrmann, The guanine cap of human guanylate-binding protein 1 is responsible for dimerization and self-activation of GTP hydrolysis. FEBS J 279, 203–210 (2012).

40. S. Ince, M. Kutsch, S. Shydlovskyi, C. Herrmann, The human guanylate-binding proteins hGBP-1 and hGBP-5 cycle between monomers and dimers only. FEBS J 284, 2284–2301 (2017).

41. S. Shydlovskyi et al., Nucleotide-dependent farnesyl switch orchestrates polymerization and membrane binding of human guanylate-binding protein 1. Proc Natl Acad Sci U S A 114, E5559–E5568 (2017).

42. A. A. Hassan, S. C. Dos Santos, V. S. Cooper, I. Sa-Correia, Comparative Evolutionary Patterns of Burkholderia cenocepacia and B. multivorans During Chronic Co-infection of a Cystic Fibrosis Patient Lung. Frontiers in microbiology 11, 574626 (2020).

43. A. A. Hassan, C. P. Coutinho, I. Sa-Correia, Burkholderia cepacia Complex Species Differ in the Frequency of Variation of the Lipopolysaccharide O antigen Expression During Cystic Fibrosis Chronic Respiratory Infection. Frontiers in cellular and infection microbiology 9, 273 (2019).

44. A. A. Hassan et al., Structure of O antigen and Hybrid Biosynthetic Locus in Burkholderia cenocepacia Clonal Variants Recovered from a Cystic Fibrosis Patient. Frontiers in microbiology 8, 1027 (2017).

45. D. Conant et al., Inference of CRISPR Edits from Sanger Trace Data. CRISPR J 5, 123–130 (2022).

46. K. Labun et al., CHOPCHOP v3: expanding the CRISPR web toolbox beyond genome editing. Nucleic acids research 47, W171–W174 (2019).

47. M. Kutsch, C. Gonzalez-Prieto, J. Coers, The GBP1 microcapsule interferes with IcsA-dependent septin cage assembly around Shigella flexneri. Pathog Dis 79, (2021).

48. J. L. Schmid-Burgk et al., Caspase-4 mediates non-canonical activation of the NLRP3 inflammasome in human myeloid cells. Eur J Immunol 45, 2911–2917 (2015).

49. C. M. Lopez, D. A. Rholl, L. A. Trunck, H. P. Schweizer, Versatile dual-technology system for markerless allele replacement in Burkholderia pseudomallei. Applied and environmental microbiology 75, 6496–6503 (2009).

50. J. M. Haldeman et al., Creation of versatile cloning platforms for transgene expression and dCas9-based epigenome editing. Nucleic acids research 47, e23 (2019).

51. S. Chen, Y. Zhou, Y. Chen, J. Gu, fastp: an ultra-fast all-in-one FASTQ preprocessor. Bioinformatics 34, i884–i890 (2018).

52. P. J. Kersey et al., Ensembl Genomes: an integrative resource for genome-scale data from non-vertebrate species. Nucleic acids research 40, D91–97 (2012).

53. A. Dobin et al., STAR: ultrafast universal RNA-seq aligner. Bioinformatics 29, 15–21 (2013).

54. M. I. Love, W. Huber, S. Anders, Moderated estimation of fold change and dispersion for RNA-seq data with DESeq2. Genome Biol 15, 550 (2014).

55. W. Huber et al., Orchestrating high-throughput genomic analysis with Bioconductor. Nat Methods 12, 115–121 (2015).

56. N. Ignatiadis, B. Klaus, J. B. Zaugg, W. Huber, Data-driven hypothesis weighting increases detection power in genome-scale multiple testing. Nat Methods 13, 577–580 (2016).

57. Y. Benjamini, Y. Hochberg, Controlling the False Discovery Rate - a Practical and Powerful Approach to Multiple Testing. J R Stat Soc B 57, 289–300 (1995).

58. M. Stephens, False discovery rates: a new deal. Biostatistics 18, 275–294 (2017).

59. R. Edgar, M. Domrachev, A. E. Lash, Gene Expression Omnibus: NCBI gene expression and hybridization array data repository. Nucleic acids research 30, 207–210 (2002).

60. M. Tamigney Kenfack et al., Deciphering minimal antigenic epitopes associated with Burkholderia pseudomallei and Burkholderia mallei lipopolysaccharide O antigens. Nat Commun 8, 115 (2017).

