## Supplementary figures and images for "Human giant GTPase GVIN1 forms an antimicrobial coatomer around the intracellular bacterial pathogen *Burkholderia thailandensis*"

### supplemental figures

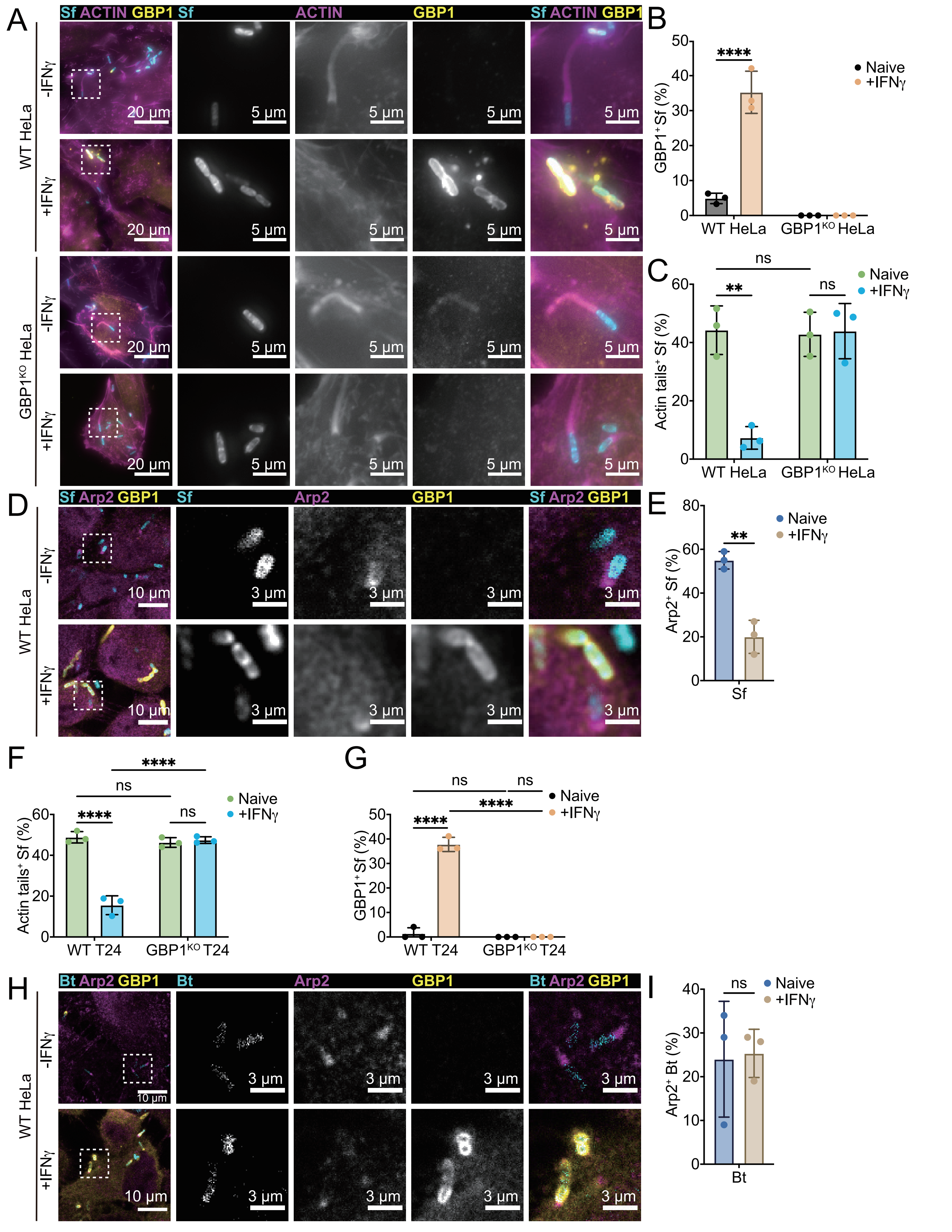

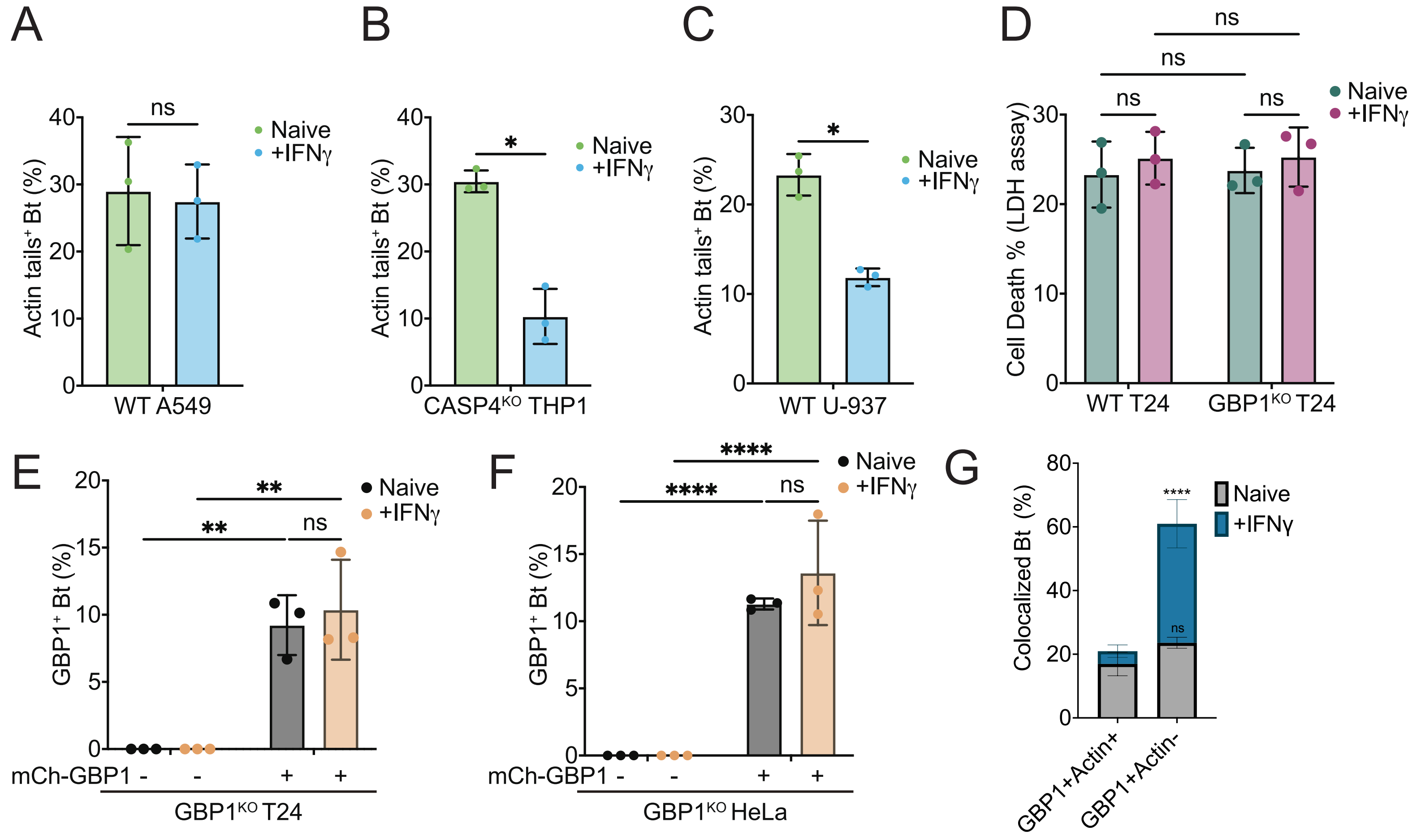

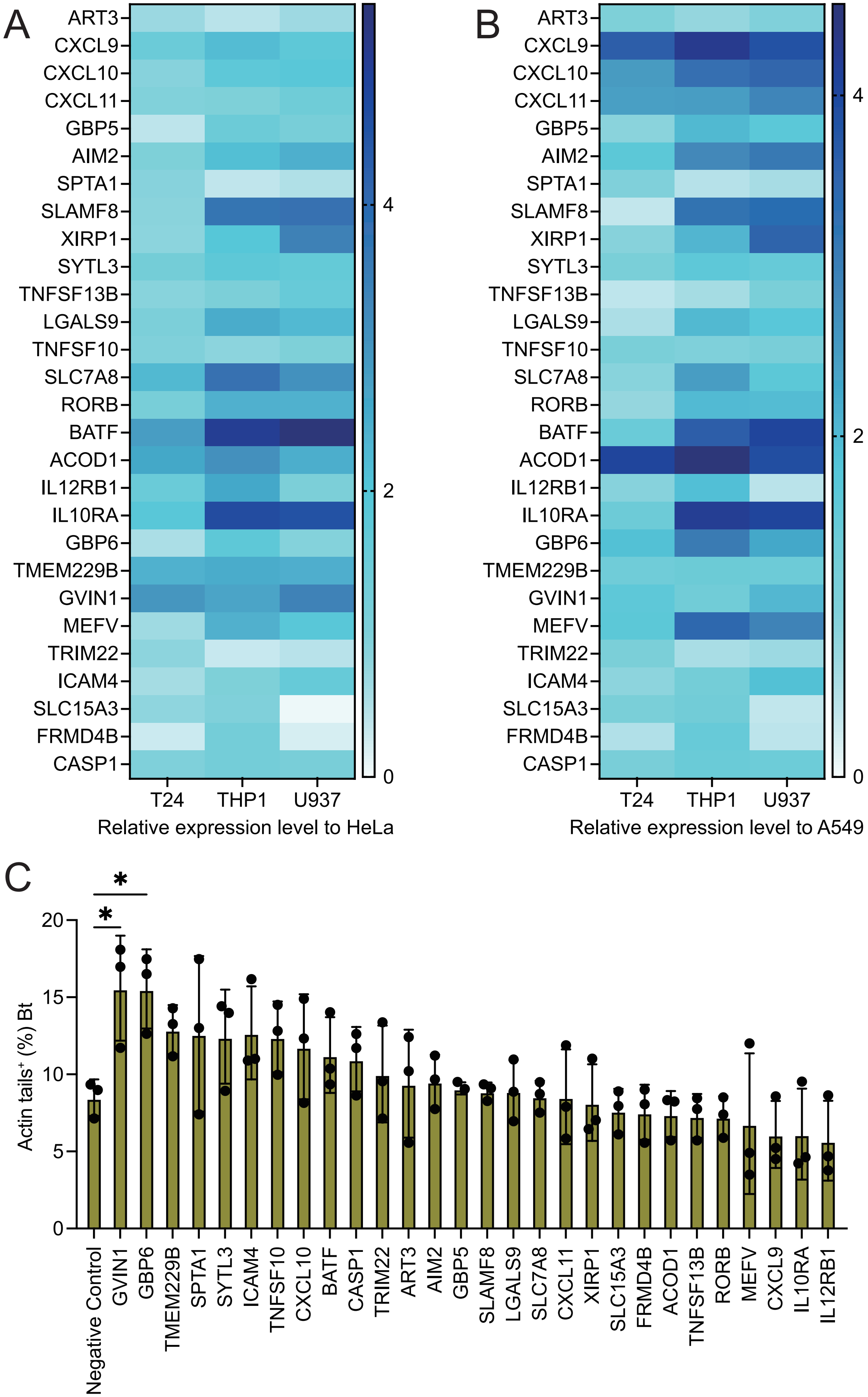

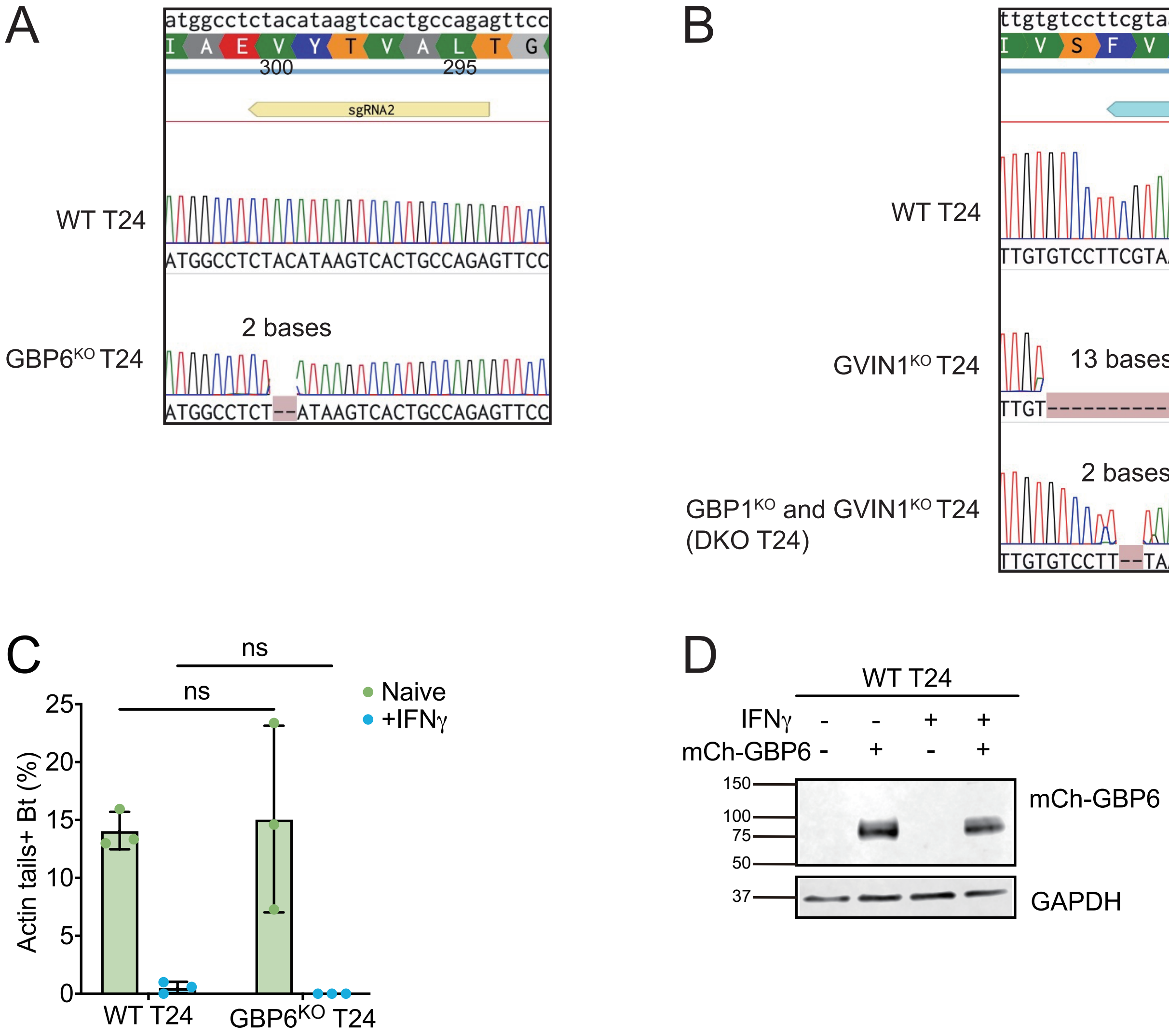

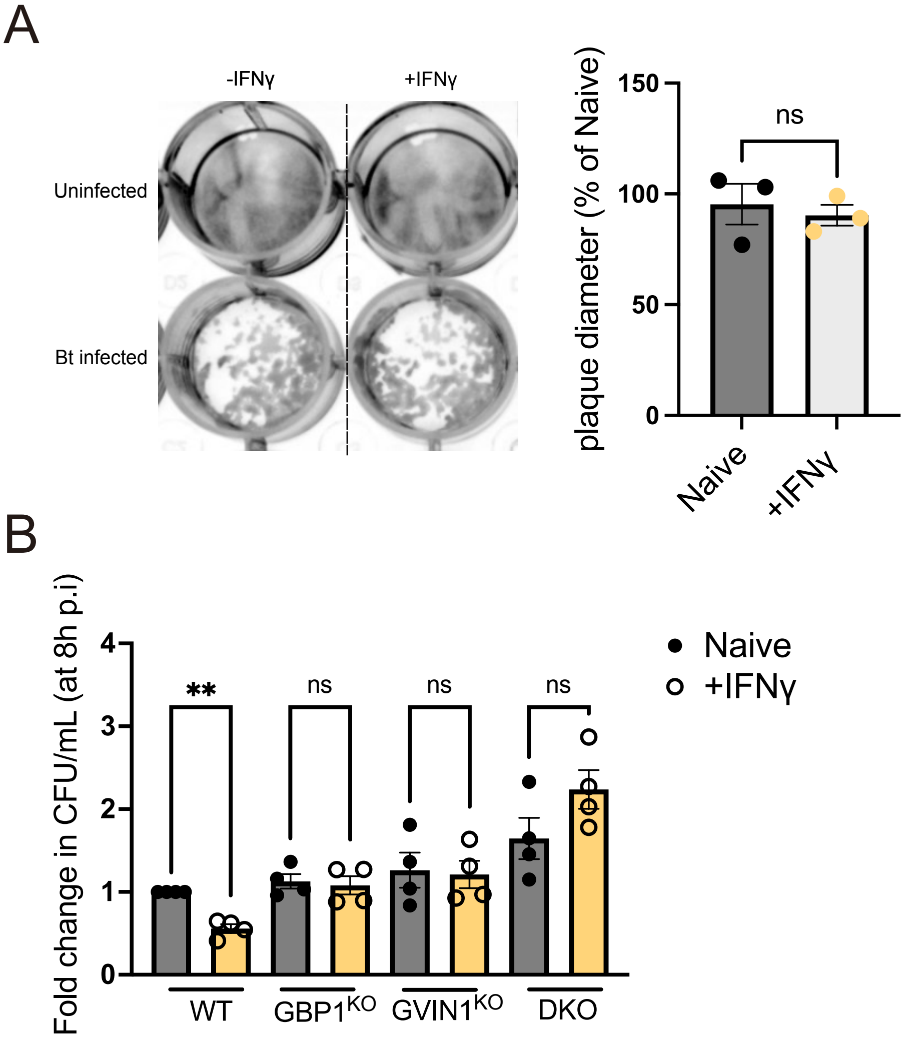

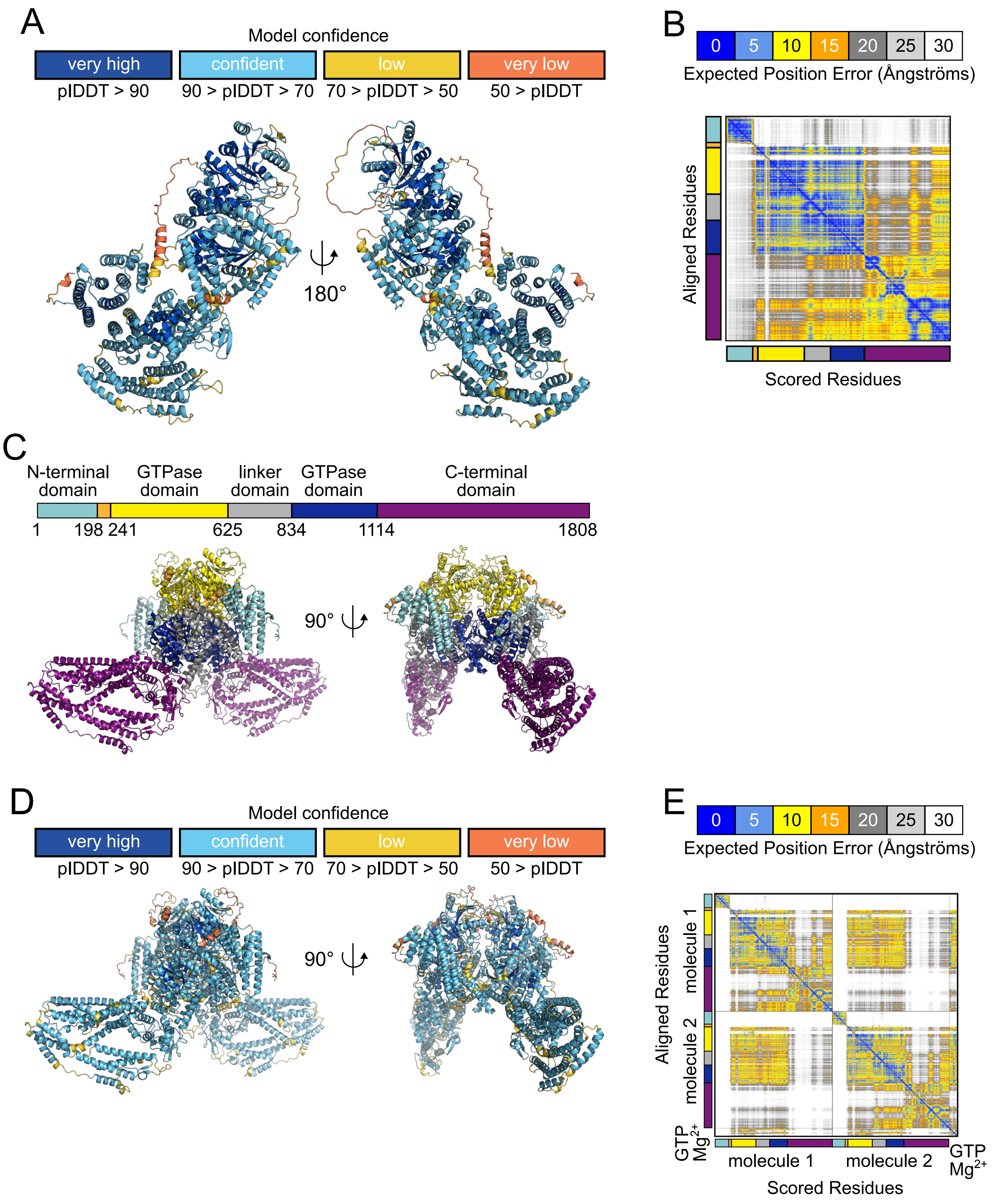

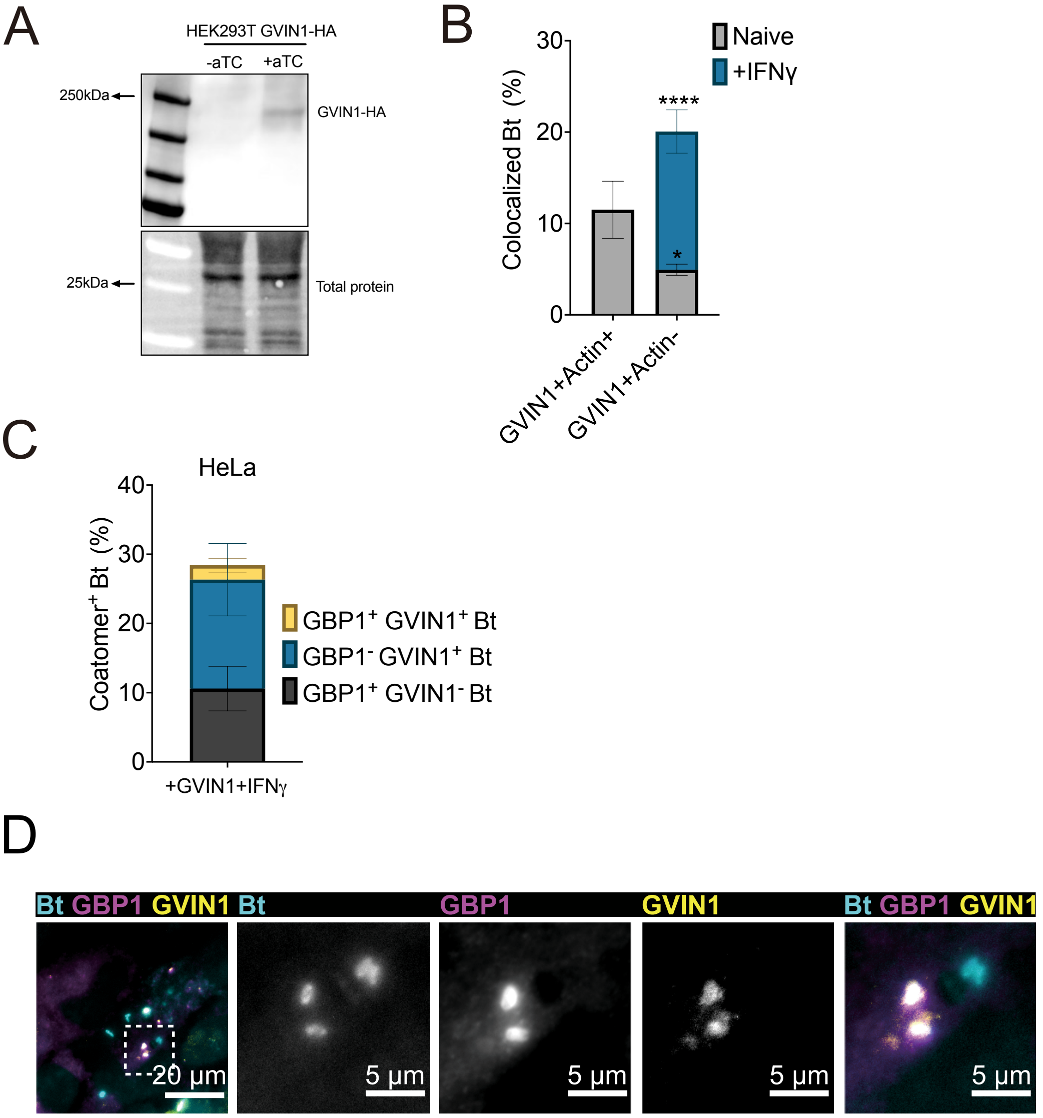

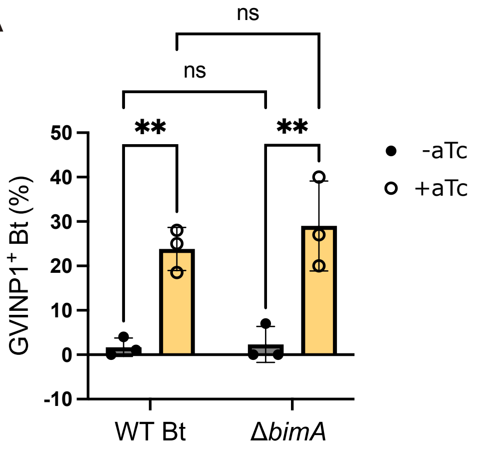

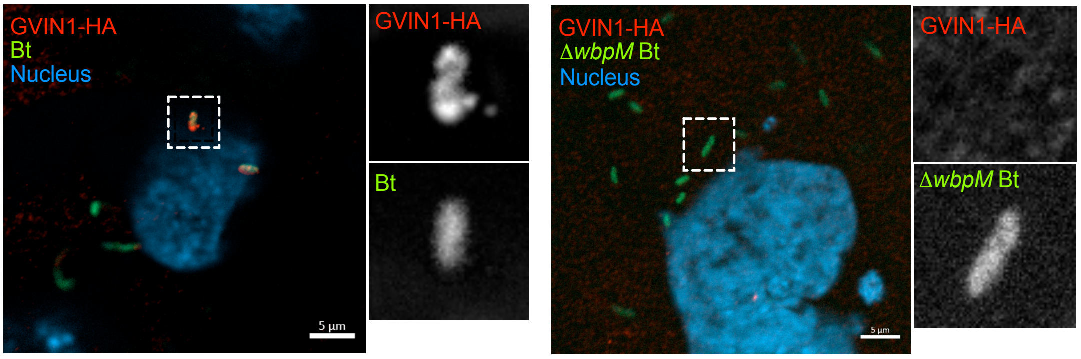

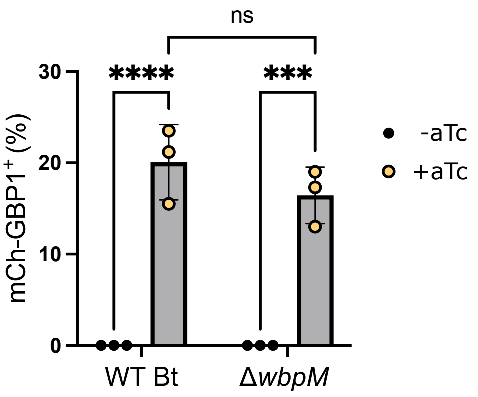

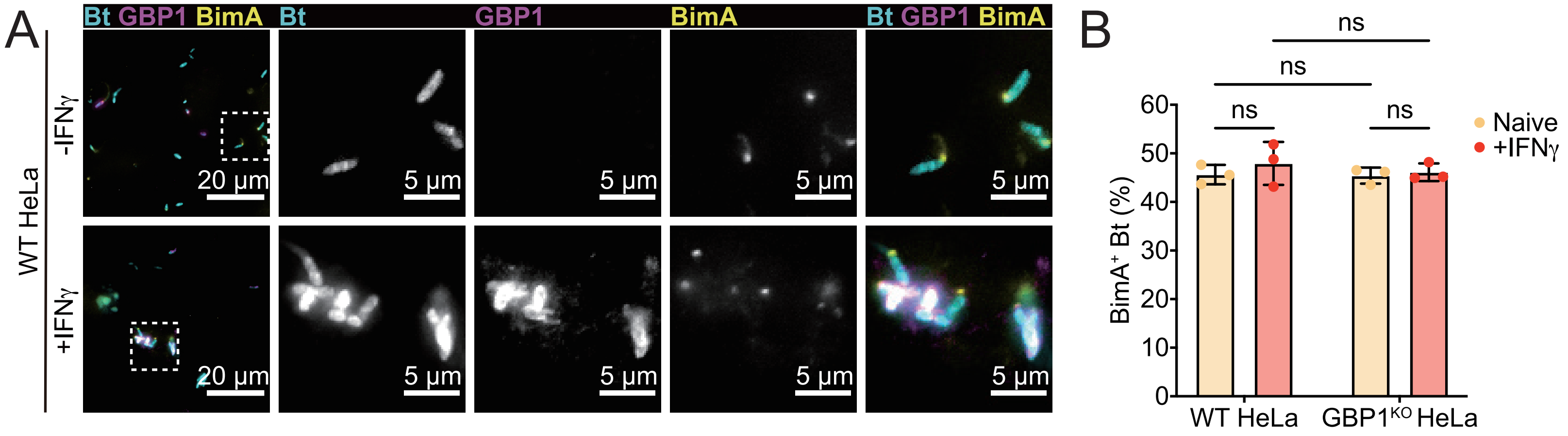
