## Supplemental figure legends for "Human giant GTPase GVIN1 forms an antimicrobial coatomer around the intracellular bacterial pathogen *Burkholderia thailandensis*"

### SUPPLEMENTARY INFORMATION

#### SUPPLEMENTARY FIGURE LEGENDS

**Fig.S1. GBP1 restrict actin tail formation by *S. flexneri* in both HeLa and T24 cells.** (A-C) Naïve or 200 U/mL IFNγ primed WT or GBP1$^{KO}$ HeLa cells were infected with *ΔipaH9.8 S. flexneri* at an MOI of 10. Cells were fixed at 2 hours and 30 minutes post infection and stained for actin and GBP1. Percentages of GBP1-positive *S. flexneri* (B) and bacteria with actin tails (C) were quantified. (D-E) Naïve or 200 U/mL IFNγ primed WT HeLa cells were infected with *ΔipaH9.8 S. flexneri* at an MOI of 10. Cells were fixed at 2 hours and 30 minutes post infection and stained for Arp2 and GBP1. Percentages of Arp2-positive *S. flexneri* (E) were quantified. (F-G) Naïve or 200 U/mL IFNγ primed WT or GBP1$^{KO}$ T24 cells were infected with *ΔipaH9.8 S. flexneri* at an MOI of 10. Cells were fixed at 2 hours and 30 minutes post infection and stained for actin and GBP1. Percentages of GBP1-positive *S. flexneri* (H) and bacteria with actin tails (I) were quantified. (H-I) Naïve or 200 U/mL IFNγ primed WT HeLa cells were infected with WT *B. thailandensis* at an MOI of 100. Cells were fixed at 8 hpi and stained for Arp2 and GBP1. Percentages of Arp2-positive *B. thailandensis* (G) were quantified. All bar graphs show 3 independent biological replicates, presented as mean ± SD. Two-way ANOVA with Tukey's multiple comparison tests were performed for (B), (C), (F), and (G); and unpaired t-test was performed for (E) and (I). Specific p-values indicated as follows: **, $p < 0.01$; ****, $p < 0.0001$; ns, not significant.

**Fig.S2. Complementary analysis of distinct phenotypes of human cell lines.** (A) Naïve or 200 U/mL IFNγ primed WT A549 cells were infected with WT *B. thailandensis* at an MOI of 100. Cells were fixed at 8 hpi and stained for actin. Percentages of *B. thailandensis* that formed actin tails(A) were quantified. (B-C) CASP4$^{KO}$ THP1 and U-937 cells were differentiated to macrophages by incubation with 50ng/mL PMA for 48 hours and were grown for another 48 hours in standard media without PMA. T24 and U-937 derived macrophages were subsequently primed with 200 U/mL IFNγ or left untreated, and then infected with WT *B. thailandensis* at an MOI of 20. Cells were fixed at 8 hpi and stained for actin. Percentages of *B. thailandensis* with actin tails (B, C) were quantified. Macrophages were more susceptible to *B.thailandensis* infection than epithelial cells, therefore infections were carried with lower MOI and shorter time. (D) Naïve or 200 U/mL IFNγ primed WT or GBP1$^{KO}$ T24 cells were infected with *B. thailandensis* at an MOI of 100. Cell death was assessed at 8 hpi via the LDH assay and is reported as the percent fluorescence relative to the lysed control. (E) Naïve or 200 U/mL IFNγ primed GBP1$^{KO}$ T24 + mCh-GBP1 cells were treated with or without 1 μg/mL aTc for overnight, and then infected with WT *B. thailandensis* at an MOI of 100. Cells were fixed at 8 hpi. Percentages of GBP1-positive *B. thailandensis* were quantified. (F) Naïve or 200 U/mL IFNγ primed GBP1$^{KO}$ HELA + mCh-GBP1 cells were treated with or without 1 μg/mL aTc for overnight, and then infected with WT *B. thailandensis* at an MOI of 100. Cells were fixed at 8 hpi. Percentages of GBP1-positive *B. thailandensis* were quantified. (G) WT T24 or T24 GBP1$^{KO}$ cells expressing mCh-GBP1 were infected with *B. thailandensis* at an MOI of 100. The percentage of mCh-GBP1 coated bacteria with or without actin tails was enumerated in naïve and IFNγ primed conditions. All bar graphs show 3 independent biological replicates, presented as mean ± SD. Two-way ANOVA with Tukey's multiple comparison tests were performed for (D), (E), (F) and (G); and unpaired t-test was performed for (A), (B) and (C). Specific p-values indicated as follows: *, $p < 0.05$; **, $p < 0.01$; ****, $p < 0.0001$; ns, not significant.

**Fig.S3 Relative transcriptional expression of 28 ISGs and their impact on GBP1-independent restriction of *B. thailandensis* actin tail formation.** (A-B) Heat maps of log2-transformed fold changes of mRNA expression levels comparing indicated IFN$\gamma$-primed cell lines with IFNγ-primedHeLa cells (A), or with IFNγ-primed A549 cells (B). A pseudocount 0.01 was added to avoid the zero-frequency problem. Listed genes are sorted by the extent to which gene expression is upregulated with IFNγ priming, from high to low. (C) GBP1$^{KO}$ T24 cells treated with siRNAs targeting genes listed on the x-axis were primed with 200 U/mL IFNγ overnight and infected with WT *B. thailandensis* at an MOI of 100. Cells were fixed at 8 hpi and phalloidin stained for actin tail detection and counting. Percentages of *B. thailandensis* with actin tails were quantified. This graph corresponds to the heatmap in Fig.4B**.** All bar graphs show 3 independent biological replicates, presented as mean ± SD. Ordinary one-way ANOVA was performed, with specific p-values indicated as follows: *, $p < 0.05$

**Fig.S4 GBP6 is dispensable for IFNγ-induced blockade in actin tail formation.** (A) Sanger sequencing confirms a frame-shift mutation in GBP6$^{KO}$ T24 cells, aligned to the wildtype genomic sequence of the targeted region. The numbers reflect the position of the amino acid in the protein sequence, starting with the start amino acid methionine of GBP6. (B) Sanger sequencing confirms frame-shift mutations in GVIN1$^{KO}$ T24 and GBP1$^{KO}$ and GVIN1$^{KO}$ DKO T24 cells, aligned to the wildtype genomic sequence of the targeted region. The numbers reflect the position of the amino acid in the predicted protein sequence, starting with the start amino acid methionine of the longest ORF of GVIN1. (C) Naïve or 200 U/mL IFNγ primed WT or GBP6$^{KO}$ T24 cells were infected with WT *B. thailandensis* at an MOI of 100. Cells were fixed at 8 hpi and stained for actin. Percentages of *B. thailandensis* with actin tails were quantified. (D) Immunoblotting for GBP6 in Naïve or 200 U/ml IFNγ primed WT T24 cells, or T24 cells with an ectopically expressed mCherry-GBP6. All bar graphs show 3 independent biological replicates, presented as mean ± SD. Two-way ANOVA with Tukey's multiple comparison tests were performed, with specific p-values indicated as follows: ns, not significant.

**Fig.S5. Effect of IFNγ priming on bacterial burden and cell-to-cell spread in T24 cells.** (A) Naïve and 200 U/mL IFNγ primed WT T24 cells were infected with *B. thailandensis* at an MOI of 0.1. At 18 hpi cells were fixed and stained with 1% crystal violet solution for 10 min followed by washing with distilled water. Plates were imaged and the plaque area was quantified with ImageJ® software. (B) Naïve or 200 U/mL IFNγ primed WT T24, T24 GBP1KO, GVIN1KO and DKO cells were infected with *B. thailandensis* at an MOI of 10. At 8 hpi host cells were lysed and lysates plated on LB agar plates to determine CFU/mL All bar graphs show at least 3 independent biological replicates, presented as mean ± SD. Unpaired t-test was performed (A) and (B) with specific p-values indicated as follows: **, $p < 0.01$; ns, not significant.

**Fig.S6 Confidence score for the GVIN1 AlphaFold model and model of GTP-bound GVIN1 dimer.** (A) pLDDT scores are shown as a measure of local confidence in the GVIN1 AlphaFold model per residue. (B) Predicted aligned error (PAE) of the relative position and orientation between the different residues in the 3D model. (C) AlphaFold model of the GTP-bound GVIN1 dimer. (D) pLDDT scores are shown as a measure of local confidence in the GVIN1 dimer model per residue. (E) Predicted aligned error (PAE) of the relative position and orientation between the different residues, ligands, and ions in the 3D model.

**Fig.S7. Detection of HA-tagged GVIN1 by Western blotting and assessment GVIN1 coatomer formation.** (A) Immunoblot showing ectopically expressed GVIN1-HA in HEK293T cells. (B) Naïve or 200 U/mL IFNγ primed HeLa cells expressing GVIN1-HA were infected with *B. thailandensis* at an MOI of 100. The percentage of GVIN1 coated bacteria with or without actin tails was enumerated (C) Naïve or 200 U/mL IFNγ primed HeLa cells ectopically expressing

GVIN1-HA were infected with *B. thailandensis* at an MOI of 100. Cells were fixed at 8 hpi and stained for endogenous GBP1 and GVIN1-HA. The magnified micrograph highlights rare incidences of bacteria coated by both GBP1 and GVIN1 (D) Percentages of *B. thailandensis* coated with GBP1, GVIN1, or both were quantified. In (B), (C) and (D). Two-way ANOVA with Tukey's multiple comparison tests were performed (B) and (D), with specific p-values indicated as follows: ****, $p < 0.0001$; ns, not significant.

**Fig.S8. Assessment of *bimA* and *wbpM* mutations on coatomer formation.** (A**)** Bar graph shows the percentage of WT and Δ*bimA B.thailandensis* encased in GVIN1 coatomers in HeLa cells. GVIN1-HA expression was induced with aTc. (B) Representative images of GVIN1-HA localization to WT and Δ*wbpM B.thailandensis* in aTc-treated HeLa cells at 8 hpi. The corresponding quantification of these data are depicted in Fig. 5E. (C) Naïve and IFNγ primed T24 GBP1$^{KO}$ cells harboring an aTc-inducible mCh-GBP1 construct were infected with WT or Δ*wbpM B.thailandensis* at an MOI of 100. Cells were fixed at 8 hpi and percentages of bacteria coated with mCh-GBP1 were quantified. All bar graphs show 3 independent biological replicates, presented as mean ± SD. Two-way ANOVA with Tukey's multiple comparison tests were performed (A) and (C), with specific p-values indicated as follows: **, $p < 0.01$; ****, $p < 0.0001$; ns, not significant.

**Fig.S9. IFNγ priming has no impact on *B. thailandensis* BimA outer membrane expression in HeLa cells.** (A-B) Naïve or 200 U/mL IFNγ primed WT or GBP1$^{KO}$ HeLa cells were infected with *B. thailandensis* expressing FLAG-tagged BimA at an MOI of 100. Cells were fixed at 8 hpi and stained for GBP1 and FLAG-BimA. Percentages of BimA-positive *B. thailandensis* (B) were quantified. All bar graphs show 3 independent biological replicates, presented as mean ± SD. Two-way ANOVA with Tukey's multiple comparison tests were performed, with specific p-values indicated as follows: ns, not significant.
