## Supplemental Table 1 for "Human giant GTPase GVIN1 forms an antimicrobial coatomer around the intracellular bacterial pathogen *Burkholderia thailandensis*"

### Table 1: List of siRNA oligomers used in ‘mini-screen’

| Gene Symbol | Oligomers Sequence |
|---|---|
| GVIN1 | UUAAAGGUCAUGUCCUCCCAU |
| GVIN1 | UUCAAUGCAUUUCAAUUCCUG |
| GVIN1 | UUAAGAUCUCAAUUAAUUGAG |
| GVIN1 | UUAUAUUUCUUAUUCACCUUG |
| GBP1 | CGAAAGGCAUGUACCAUAA |
| GBP1 | GAACAGGAGCAACUACUAA |
| GBP1 | CAGAUGAGUACCUGACAUA |
| GBP1 | CUGAGAAGAUGGAGAACGA |
| SLC7A8 | GGACAGAGGAGGCUAAUGA |
| SLC7A8 | UCAACUACCUCUUCUAUGG |
| SLC7A8 | CCACGAAGGACAAGGACGU |
| SLC7A8 | UGGCCAUGAUCCACGUGAA |
| IL12RB1 | GGAGGUCACUUACCGACUA |
| IL12RB1 | GGAUAUCCAGUGAUCGUUA |
| IL12RB1 | ACAUCAAGGUGUCCAAGUU |
| IL12RB1 | CGCCAUAUCCGGAUGCAGA |
| TNFSF13B | GCACAAAUAUCACUGGAUG |
| TNFSF13B | GAAAUAAGCGUGCCGUUCA |
| TNFSF13B | AAAUAUGCCUGAAACACUA |
| TNFSF13B | GAUAAGACCUACGCCAUGG |
| ACOD1 | UCGACAAAGUGCCGAGAAG |
| ACOD1 | CCCUAUUUGUCAAGCCGAA |
| ACOD1 | GGACGUGGCCUUUAAGCGU |
| ACOD1 | UGAAGUGCAAGGCCGAUUA |
| MEFV | GACCACUCCUCAAGAGAUA |
| MEFV | GAGAAUGGCUACUGGGUGG |
| MEFV | GCCCGCAAAUCCAGAAAUU |
| MEFV | GCAUAUGACACCCGCGUAU |
| ICAM4 | GCAAGUGCCUAGCUAUGAA |
| ICAM4 | GGAAGUCAGUGCAGCUCAA |
| ICAM4 | GGAAGCCGGGUCAUCUAUU |
| ICAM4 | GCGCAAAGCCCCAAGGGUA |
| IL10RA | GAAAGUACCUGCUAUGAAG |
| IL10RA | GAAACAGGAUCCUCUAGAA |
| IL10RA | GGACACCCAUCCCAAAUCA |
| IL10RA | GAAGAGAAGGCAACCAAGA |
| SLC15A3 | CAAGUGUCCUCUAUGCUUA |
| SLC15A3 | CGUCACGGCUCUCCUAUUU |
| SLC15A3 | GAAGAUCUUGCCCGUCAUG |
| SLC15A3 | GGUGGCAGAUCCCUCAGUA |
| FRMD4B | CCAUAGGGCUGUCGGAUUA |
| FRMD4B | CAAUAAACGAAUACCGAAU |
| FRMD4B | AAAUCGAAGUAGAGAGCGA |
| FRMD4B | CGGAGAUGGUACACGGAGA |
| TNFSF10 | UGGCAUUGCUUGUUUCUUA |
| TNFSF10 | AGAAAUAGUUGUUGGUCUA |
| TNFSF10 | GACCCUAUAUUGUUGAUGA |
| TNFSF10 | ACAAACAAAUGGUCCAAUA |
| TRIM22 | GUACGCACCUGCACAUUUA |
| TRIM22 | CACCAAACAUUCCGCAUAA |
| TRIM22 | CCAGAUAUAGACCUCAAUA |
| TRIM22 | AGAAUUAUAUCCAGAUCGA |
| CASP1 | GACUCAUUGAACAUAUGCA |
| CASP1 | AGACAUCCCACAAUGGGCU |
| CASP1 | GAAUAUGCCUGUUCCUGUG |
| CASP1 | CCGCAAGGUUCGAUUUUCA |
| CXCL9 | GAAUGGAGUUCAAACAUGU |
| CXCL9 | CCAAGGGACUAUCCACCUA |
| CXCL9 | AAACCCAGAUUCAGCAGAU |
| CXCL9 | GUAAAACACUUGCGGAUAU |
| GBP5 | GGAAAUAGAUGGGCAACUU |
| GBP5 | UCUGAGAGAUUUCUGCUUA |
| GBP5 | GCCCAGAGGACUGUUAAUA |
| GBP5 | AUGAGCAGCUGAAGGUUAA |
| BATF | GUACAGCGCCCACGCAUUC |

| | |
|---|---|
| BATF | GAAACAGAACGCGGCUCUA |
| BATF | GAACGCGGCUCUACGCAAG |
| BATF | AGAGUUCAGAGGAGGGAGA |
| ART3 | CCAAAUACCUGAAGAUAAA |
| ART3 | GGAAAUAUCAACAAUCCUA |
| ART3 | CUACAACCCUGGUGAGAAA |
| ART3 | UGUUAGACAUGGCAGAUAA |
| SLAMF8 | CUACUUCCCAUUACAGUUA |
| SLAMF8 | GGGAAGGCCUCCUACAAAG |
| SLAMF8 | CGAAAUAACCUAUAGCUGG |
| SLAMF8 | GUUCAUUGCUGUAGAAAGG |
| SPTA1 | GGACAAAGCUUGGAGACUA |
| SPTA1 | GCACCGAAGUUCUGCAUAA |
| SPTA1 | GCAGAGGGCAAGUCAUAUA |
| SPTA1 | GAACCACGCAUUCAAGAGA |
| AIM2 | GAAACGAGGACACAAUGAA |
| AIM2 | UCAGACGAGUUUAAUAUUG |
| AIM2 | GGAUAGGUUUAAGUUCUUU |
| AIM2 | GUUCAUAGCACCAUAAAGG |
| SYTL3 | GUACAGAGAUGGGCAAUUU |
| SYTL3 | GUAAGAACCUUGCCUAUGG |
| SYTL3 | GAAGACAGCGUUCCUCAGA |
| SYTL3 | GGUGGAAAGGAGCGAAGAA |
| XIRP1 | GAAGACGGCCUGUGAGAUA |
| XIRP1 | GCAGGUGUCUCGUCAGAAA |
| XIRP1 | GGAGACAUCAAGAGAGUAU |
| XIRP1 | GAGAGCAUCAUCCAUGUUC |
| LGALS9 | GAGGGUACGUGGUGUGCAA |
| LGALS9 | GCUCAGAUUUCAAGGUGAU |
| LGALS9 | GGUCAGCACCUGUUUGAAU |
| LGALS9 | GCAACACCCAGAUCGACAA |
| CXCL10 | GAGAAGAGAUGUCUGAAUC |
| CXCL10 | AAUCGAAGGCCAUCAAGAA |
| CXCL10 | AAUCCAAGGUCUUUAGAAA |
| CXCL10 | UGAAAGCAGUUAGCAAGGA |
| CXCL11 | GCUGUGAUAUUGUGUGCUA |
| CXCL11 | GCAGUGAAAGUGGCAGAUA |
| CXCL11 | GCCUAAAUCCCAAAUCGAA |
| CXCL11 | AGUGAUUAUUACCCUGAAA |
| GBP6 | CCGAUGAGCUAACUGCAAU |
| GBP6 | GGACUAAUGGAAAGUAUCU |
| GBP6 | CAACAGAGCCUGGGCGACA |
| GBP6 | CAUGGAGAGUCAUCGUAAA |
| TMEM229B | GGACCUACCUGUGGGAGUU |
| TMEM229B | GCUGGUACCUGUAUGCCAU |
| TMEM229B | GCUUCGACAAGGACGCUGA |
| TMEM229B | CAUUACAAAGUGACGGUAU |
| RORB | GAUCAAAUUCUACUUCUGA |
| RORB | UCAAACAGAUAAAGCAAGA |
| RORB | CCAAGUCUGAGGGUUAUUA |
| RORB | GCGGAUAACAGGCUUCAUG |
